# Dysregulation of splicing-related proteins in prostate cancer is controlled by FOXA1

**DOI:** 10.1101/509034

**Authors:** John G. Foster, Rebecca Arkell, Marco Del Giudice, Chinedu Anene, Andrea Lauria, John D. Kelly, Nicholas R. Lemoine, Salvatore Oliviero, Matteo Cereda, Prabhakar Rajan

## Abstract

Prostate cancer (PCa) is genomically driven by dysregulation of transcriptional networks involving the transcriptional factors (TFs) FOXA1, ERG, AR, and HOXB13. However, the role of these specific TFs in the regulation of alternative pre-mRNA splicing (AS), which is a proven therapeutic vulnerability for cancers driven by the TF MYC, is not described. Using transcriptomic datasets from PCa patients, we tested for an association between expression of *FOXA1, ERG, AR, HOXB13*, and *MYC*, and genes involved in AS - termed splicing-related proteins (SRPs), which have pleiotropic roles in RNA metabolism. We identified FOXA1 as the strongest predictor of dysregulated SRP gene expression, which was associated with PCa disease relapse after treatment. Subsequently, we selected a subset of FOXA1-binding and actively-transcribed SRP genes that phenocopy the FOXA1 dependency of PCa cells, and confirmed *in vitro* via knockdown and over-expression that FOXA1 regulates SRP gene expression. Finally, we demonstrated the persistence of a FOXA1-SRP gene association in treatment-relapsed castration-resistant PCa (CRPCa) patients. Our data demonstrate, for the first time, that FOXA1 controls dysregulated SRP gene expression, which is associated with poor PCa patient outcomes. Analogous to MYC-driven cancers, our findings implicate the therapeutic targeting of SRPs and AS in FOXA1-overexpressing PCa.

## Introduction

Prostate cancer (PCa) is the commonest male gender-specific cancer ^1^. Genomic characterisation of primary PCa has uncovered several molecular subtypes ^2-4^, characterised by alterations in genes encoding the transcription factors (TFs) FOXA1, ERG, and AR, which is an existing therapeutic target ^5^. The gene encoding the AR-interacting TF HOXB13 has been identified as a candidate PCa susceptibility gene ^6^. Co-operatively, these TFs reprogram the AR-associated cistrome in prostate tumourigenesis ^7-9^. Additionally, PCa susceptibility loci identified from genome-wide association studies (GWAS) fall within the cistromes of these TFs themselves ^10^, thereby demonstrating widespread dysregulation of transcriptional networks in PCa.

Recently, widespread genomic and transcriptional dysregulation of genes encoding RNA-binding proteins (RBPs) have been reported across several cancers ^11-13^. RBPs are a family of proteins with pleiotropic roles in RNA metabolism including alternative pre-mRNA splicing (AS) ^14^. Through the regulation of their target mRNAs, RBPs are associated with different oncogenic processes and patient outcomes ^15^. In PCa and other malignancies, mutations in genes coding for RBPs and changes in their expression levels have been observed ^11,16^. Moreover, AS appears to represent a cancer therapeutic vulnerability in leukaemia and breast cancer driven by the oncogenic TF MYC ^17-19^. MYC has also been shown to transcriptionally regulate expression of AS-associated RBPs in lymphoma, lung cancer and glioma pre-clinical models ^20-22^. However, little is known of the mechanisms of transcriptional dysregulation of RBP expression in PCa.

Here, we show that genes encoding RBPs and other proteins involved in AS (referred to, hereafter, as splicing-related proteins; SRPs) are globally dysregulated in PCa, and identify the TF FOXA1 as a key regulator of SRP gene expression.

## Results

### Dysregulated SRP gene expression is associated with FOXA1 in primary human PCa

We hypothesised that dysregulation of SRP genes and others involved in gene expression processes in PCa is transcriptionally controlled by one or more of the TFs FOXA1, ERG, AR or HOXB13. To test this hypothesis, we utilised published RNA-Seq gene expression data of primary untreated prostate tumours (n=409) included in The Cancer Genome Atlas (TCGA) ^2^. Transcriptomes were analysed based on expression levels (*i.e*. transcript per million reads, TPMs) of *FOXA1, ERG, AR* and *HOXB13* genes (Fig. 1A). We included *MYC* as a positive control as it is implicated in the regulation of SRP expression ^20-22^. Samples were stratified for expression of genes encoding these five TFs with a cut-off of the top 25% of gene expression by TPM defining high expression (HE) and the remainder as Rest (Fig. 1A and B, and Supplementary Data 1) ^23^.

**Fig. 1:**
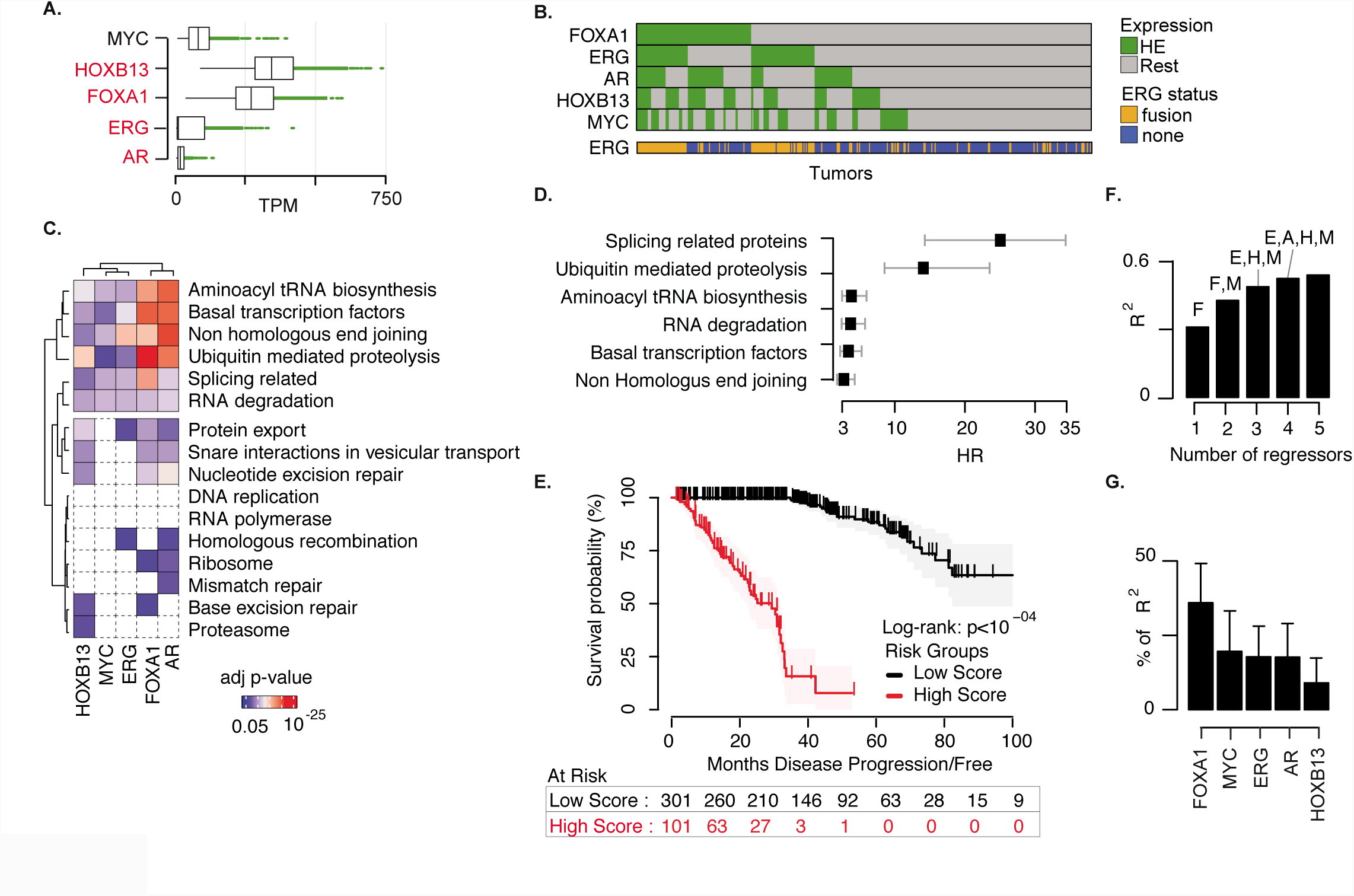
FOXA1 expression is independently associated with SRP expression in primary PCa. (A) Boxplot distributions of normalized gene expression levels (transcript per million reads, TPMs) for each of the five TF genes in primary PCa samples. Green lines and dots refer to samples with TF expression greater than the 75th percentile of its distribution across samples. PCa-related TFs are reported in red. (B) Heat map showing the stratification of tumour samples accordingly to the expression levels of each of the TF genes. Green cells depict samples exhibiting the high expression (HE) of the corresponding TF. The last row reports samples undergoing ERG gene fusion. (C) Results of the gene set analysis (GSA) for 16 biological KEGG processes related to the HE of each TF. The altered processes are hierarchically clustered on the basis of their statistical significance (i.e. adjusted p-value). Non-statistically significant processes are depicted in white dashed boxes. The side panel reports the corresponding KEGG pathway category. (D) Hazard Ratios (HR) from univariable Cox proportional hazards (PH) model using gene set scores for the six biological processes associated with FOXA1 HE. (E) Kaplan-Meier plot of survival probabilities over time for patients stratified on the 75th percentile of the SRP gene set score distribution with log rank test p-value. (F) Coefficient of determination (R2) of the linear regression model of the SRP gene expression for increasing number of regressors in the model (i.e. the TFs included in the model). F=FOXA1; M=MYC; E=ERG; H=HOXB13; A=AR. (G) Relative importance of each predictor (i.e. TF) to the R2 measured by the linear regression model including all TFs.

To determine the biological processes that are altered upon HE of the TFs, we performed a gene set analysis (GSA) (see Methods). For each TF, we compared the cumulative TPM values of genes in 16 gene sets representing Genetic Information Processing pathways accordingly to the Kyoto Encyclopedia of Genes and Genomes (KEGG) ^24^ between samples with HE of the TF gene and Rest (Fig. 1C and Supplementary Fig. 1). In doing so, GSA identified associations between expression of all TF genes and six different KEGG pathways, including the SRP gene set (Fig. 1C, Supplementary Table 1). To evaluate the impact of altered expression of genes in the six gene sets on PCa disease progression, we performed a survival analysis using Cox proportional hazards (PH) models (see Methods). Of the genes within the six gene sets, we found that dysregulated SRP gene expression showed the strongest association with disease recurrence (HR= 25.5; 95% CI=14.6-44.5) (Fig. 1D and Supplementary Table 2). Additionally, using a SRP gene set score to stratify patients (see Methods), we observed a statistically significant difference in time to disease progression between patients with a high score as compared to those with a low score (p-value < 0.0001) (Fig. 1E), thereby highlighting the importance of SRP genes in the PCa disease phenotype.

To determine which of the five TFs may be most important for SRP gene regulation, we employed a linear regression modelling approach (See Methods). We found that the overexpression of *FOXA1* gave the best results in terms of determination coefficient (R^2^=0.3, Fig. 1F) when modelling SRP gene expression using only one TF gene. Increasing the model complexity led to a closer fitting between TF overexpression and SRP gene expression levels, with the five variables giving the highest fitting (R^2^=0.54, Fig. 1F). We next measured the relative importance of each regressor in the linear model with all five TF genes using the averaging over ordering method ^25^ (see Methods). We found that *FOXA1* expression is the most important regressor contributing to 36% of the fitting of the model (Fig. 1G). Collectively, these findings suggest that, among all tested TFs, the expression of FOXA1 shows the strongest correlation with the modulation of SRP gene expression in PCa.

### The FOXA1 cistrome includes a subset of actively-transcribed SRP genes

To identify SRP candidate genes regulated by FOXA1, we performed differential expression analysis using three distinct approaches (see Methods): Firstly, we identified a total of 76 SRP genes that were significantly up-(n=54) or down-(n=22) regulated in samples with *FOXA1* HE as compared with *FOXA1* Rest (Supplementary Data 2 and Supplementary Fig. 2A). Secondly, we determined which of the 76 SRP genes had enrichment for the five TF binding sites within their promoter regions and gene bodies using the ReMap database ^26^ (Fig. 2A). Finally, we selected sites of active transcription by overlapping TF binding sites with the H3K27ac and H3K4me3 signatures. FOXA1 binding sites were most enriched within these genes compared to the other transcription factors (Fig. 2B), with 63/76 SRP genes containing FOXA1-binding sites (Supplementary Fig. 2A). Of the 63 SRP genes with FOXA1 binding sites, the majority (47/63) were up-regulated. These data suggest that FOXA1 might directly control expression of up-regulated SRP genes in PCa.

**Fig. 2:**
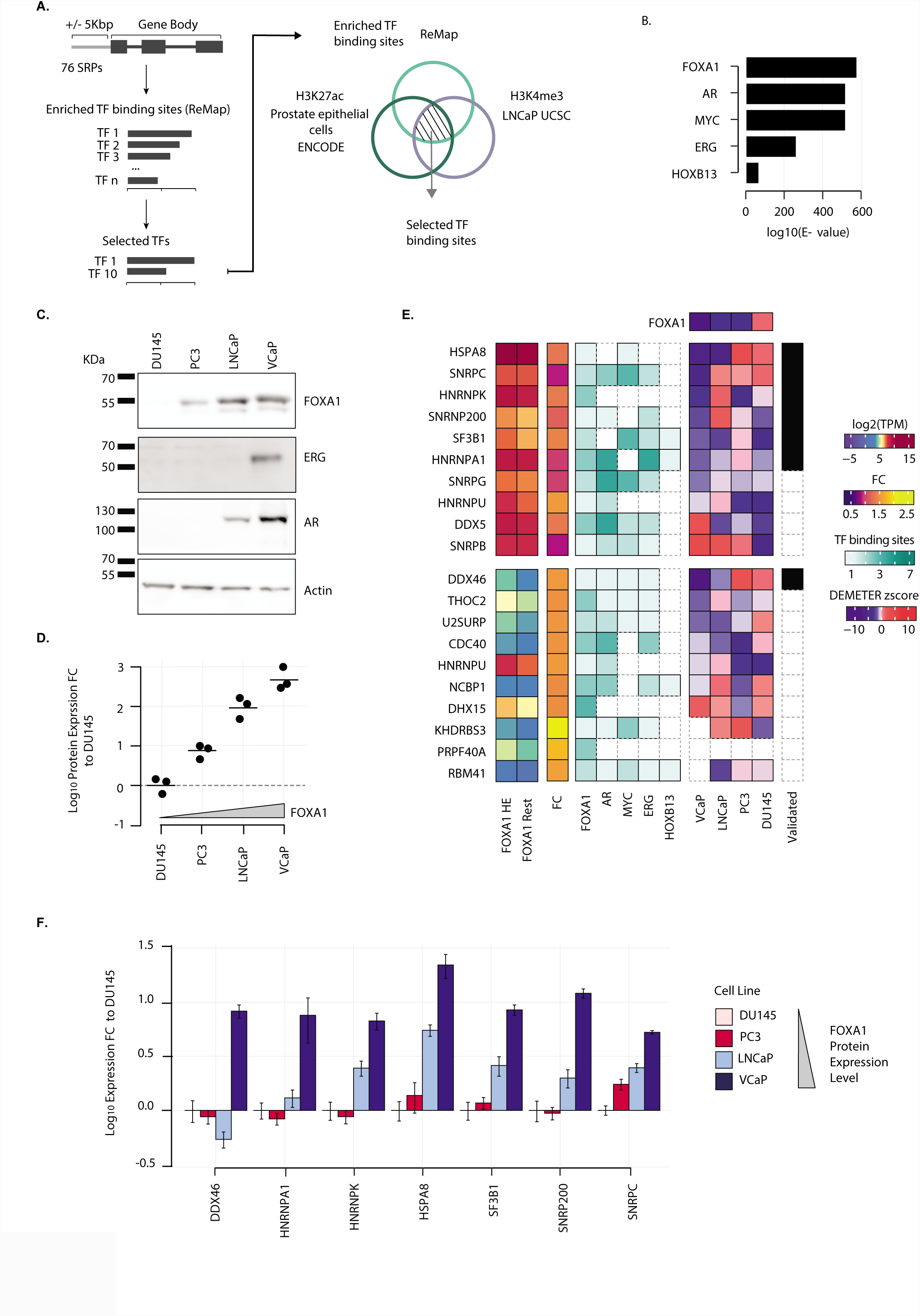
FOXA1 and SRP gene expression correlate in PCa cell line models. (A) Enrichment analysis of TF binding sites within SRP genes was performed using chromatin immunoprecipitation sequencing (ChIP-Seq) data from the ReMap database and annotation tool, as described in the Materials and Methods section. (B) Enrichment values for the five selected TFs generated from the ReMap analysis. Red indicates TFs that are significantly up-regulated in FOXA1 HE compared to Rest. (C) Representative Western blotting images of whole cell lysates from PCa cell lines using antibodies to AR, ERG and FOXA1, and actin. (D) Densitometric band quantitation was performed to calculate log_10_ relative normalized fold change (FC) in expression relative to DU145. Data from at least three independent experiments were used to calculate the means ± SEM. (E) Heatmap depicts TPM values of candidate SRPs in FOXA1 HE and rest PCa samples (first block), fold change of expression (second block), number of TF binding sites in region of active transcription (third block), DETEMER Z-score for SRP dependency in VCaP, LNCaP, PC3 and DU145 cell lines (forth block). DEMETER Z-score for FOXA1 in the four cell lines is reported. Selected SRPs for *in vitro* validation in PC cell lines are reported in black. Missing values or zeros are depicted by white dashed blocks. In the upper block SRPs are sorted in decreasing order of TPM values in FOXA1 samples. In the lower block SRPs are sorted in decreasing order of FC. (F) qRT-PCR was performed on cDNAs from different PCa cells lines, and levels of SRP transcript expression were normalized to a geometric mean of *ACTB* and *B2M* levels to calculate log_10_ relative normalized FC in expression compared to DU145 cells. Data from at least three independent experiments were used to calculate the means ± SEM.

### The expression and function of FOXA1 and SRP genes are similar in human PCa cell line models

To characterise human PCa cell line models for downstream validation, we profiled *FOXA1* expression by qRT-PCR and western blotting in DU145, PC3, LNCaP and VCaP cells (Fig. 2C and Supplementary Fig. 2B). We identified the highest level of FOXA1 expression in the AR-and ERG-positive VCaP cells, and the lowest level of expression in DU145 cells (Fig. 2D). FOXA1 has been identified as an essential PCa gene in a RNAi genome-wide loss of function screen ^27^, and in this dataset VCaP and DU145 cells harboured the greatest and least dependency (DEMETER scores) on FOXA1, respectively (Fig. 2E). To confirm this observation, we used two independent siRNA duplexes (Supplementary Table 3) to deplete VCaP and DU145 cells of FOXA1 protein (Supplementary Fig. 2C, upper panel). Following FOXA1 depletion, we observed a statistically significant reduction in cell growth in VCaP cells as compared with NSI controls, but no statistically significant change in cell growth in DU145 cells (Supplementary Fig. 2C, lower panel).

We hypothesised that FOXA1-overexpressing VCaP cells may also be dependent on FOXA1-associated SRP genes. Of the 47 up-regulated SRP genes with FOXA1-binding sites identified from the TCGA analysis (Supplementary Fig. 2A), we selected the top 10 ranked SRP genes by TPM and top 10 ranked by fold change of median TPM between FOXA1 HE samples and Rest for further analysis (Fig. 2E). Using the RNAi dataset ^27^, we ranked these 20 SRP genes by VCaP cell dependency DEMETER score, and selected seven candidate SRP genes with a DEMETER score smaller than −1 for qRT-PCR validation in four different PCa cell lines. We profiled expression of the seven candidate SRP genes in the four PCa cell lines by qRT-PCR and observed the highest levels of expression of SRP genes in FOXA1-overpressing VCaP cells as compared with DU145 cells (Fig. 2F and Supplementary Table 4). Taken together, our data identify a subset of seven up-regulated and FOXA1-associated SRP genes in primary PCa that phenocopy FOXA1 in PCa cells that over-express FOXA1.

### FOXA1 regulates SRP gene expression in vitro in human PCa cell line models

To determine whether the above candidate SRPs are regulated by FOXA1 *in vitro*, we utilized two independent siRNA duplexes to deplete PCa cell lines of FOXA1 (Fig. 3A-D, left panels and Supplementary Table 5). siRNA-mediated depletion of FOXA1 protein in VCaP cells resulted in a reduction in expression of six out of seven SRPs genes by qRT-PCR, where statistical significance was observed with at least one siRNA duplex (p-value <0.05, Fig. 3A, right panel, Supplementary Tables 6-7). A similar effect of FOXA1 depletion on SRP gene expression was observed in LNCaP (Fig. 3B, right panel, 3/7 SRPs), PC3 (Fig. 3C, right panel, 4/7 SRPs) and DU145 (Fig. 3D right, 4/7 SRPs). We then tested whether ectopic expression of FOXA1 in PC3 cells, which harbour lower FOXA1 protein level (Fig. 2C and D), would conversely result in an increase in SRP gene expression (Fig. 3E). We observed a general increase in SRP gene expression, with four SRP genes reaching statistical significance (Fig. 3E, right panel).

**Fig. 3:**
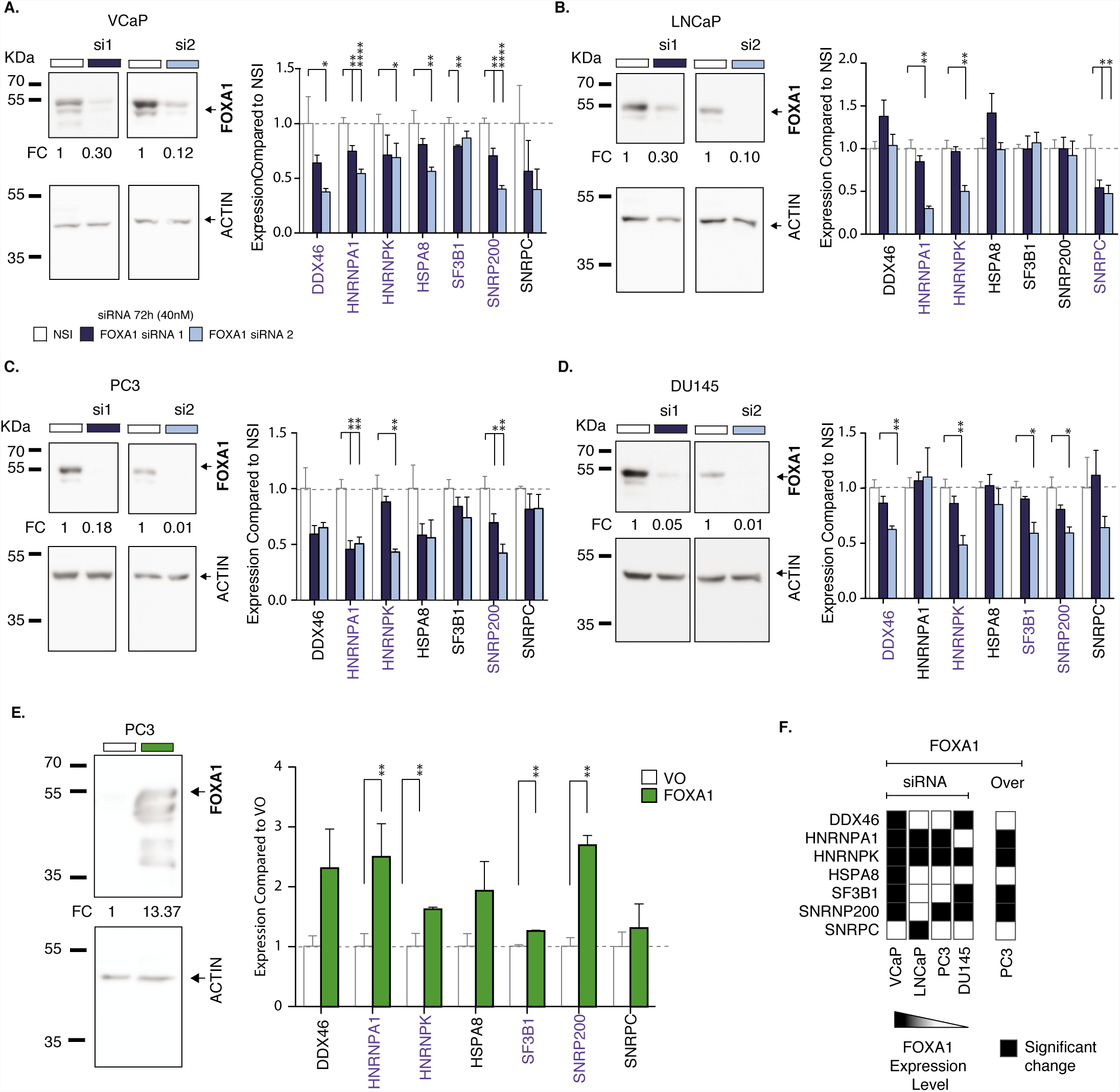
FOXA1 regulates SRP expression in PCa cell lines *in vitro*. (A) DU145, (B) PC3, (C) LNCaP, and (D) VCaP were transfected with two siRNA duplex sequences to FOXA1 (FOXA1 si1 or si2), or non-silencing (NSI) control to final concentration of 40 nM. (A-D, Left Panels) After 72 h, total cell lysates were harvested and subjected to western blotting with antibodies to FOXA1 and actin. Western blotting images shown are representative of three independent experiments, from which densitometric band quantitation was performed to calculate the mean relative normalized fold change (FC) in protein expression (shown below FOXA1 blot images). (A-D, Right Panels) qRT-PCR was performed on cDNAs and levels of SRP transcript expression were normalized to a geometric mean of *ACTB* and *B2M* levels to calculate relative normalised FC in expression compared to NSI. Data from at least three independent experiments were used to calculate the means ± SEM. (E) PC3 cells were transfected with expression vectors for pcDNA3.1-FOXA1 or vector only (VO) control (1 μg) as indicated. After 72 h, total cell lysates were harvested and subjected to western blotting with antibodies to FOXA1 and actin (E, left panel). Western blotting images shown are representative of three independent experiments, from which densitometric band quantitation was performed to calculate the mean relative normalized FC in protein expression (shown below FOXA1 blot images). (E, right panel) qRT-PCR was performed on cDNAs and levels of SRP transcript expression were normalized to a geometric mean of ACTB and B2M levels to calculate relative normalised FC in expression compared to VO. Data from at least three independent experiments were used to calculate the means ± SEM. Unpaired two-tailed T-test was used to compared groups: *p-value <0.05, **p-value <0.01, ***p-value <0.001, ****p-value <0.0001. (F) Heatmap showing significant expression changes of SRP genes (black cells) in the different PCa cell lines upon FOXA1 silencing and overexpression.

Overall, there was a statistically-significant reduction in *HNRNPK* expression levels of in all four cell lines upon FOXA1 depletion, and a statistically-significant increase upon the overexpression of FOXA1 in PC3 cells (Fig. 3F). With the exception of the DU145 cell line that expresses the lowest level of FOXA1, we observed a consistent impact of *FOXA1* depletion and overexpression on *HNRNPA1* and *HNRNPK* expression (Fig. 3F). These data demonstrate that FOXA1 can regulate SRP gene expression in PCa cell lines, with the most consistent effect observed in FOXA1-overexpressing VCaP cells.

### ERG and AR do not appear to regulate SRP gene expression in human PCa cell line models

Of the candidate SRP genes, a number were bound by AR and ERG as well as FOXA1 (Fig. 2E). Since FOXA1 has been implicated as a pioneer factor ^28^ and is co-opted by AR ^29^ and ERG ^30^ in PCa cells, we hypothesised that the observed FOXA1 regulated SRP gene expression is mediated by AR and ERG. To test this hypothesis, we used siRNA to deplete ERG in the VCaP cells, and AR in VCaP and LNCaP cells (Supplementary Fig. 3, left panels). Surprisingly, following siRNA-mediated ERG knockdown, we observed an increase in expression of two out of seven SRP genes by qRT-PCR (Supplementary Fig. 3A, right panel), which appeared to be due to an increase in FOXA1 protein expression as determined by western blotting (Supplementary Fig. 3A, left panel). siRNA-mediated AR knockdown in LNCaP and VCaP cells did not significantly impact on FOXA1 protein expression nor SRP gene expression (Supplementary Fig. 3B and C). Consistent with our findings *in silico* (Fig. 1), these data confirm that AR and ERG are not as important as FOXA1 in SRP gene regulation in PCa, and suggest that this role may be a novel AR-independent function of FOXA1.

### FOXA1-associated SRP gene dysregulation is persistent in advanced treatment-relapsed PCa

We sought to determine whether the dysregulation of SRP genes persists in late-stage, metastatic, castration-resistant PCa (CRPCa) which inevitably develops following longstanding androgen deprivation therapy (ADT) ^5^, and whether this is associated with expression of *FOXA1, ERG, AR, HOXB13* and *MYC*. We utilised published RNA-Seq gene expression data of CRPCa samples from patients (n=118) included in the Stand Up to Cancer (SU2C) study ^31^. As before, transcriptomes were stratified by expression levels (*i.e*. TPMs) of TFs (Fig. 4A) with a cut-off of the top 25% of gene expression by TPM defining HE. We did not detect a statistically significant difference in the levels of *FOXA1* expression between the *FOXA1* HE tumours in the TCGA and SU2C datasets (two-tailed Wilcoxon rank sum test, Supplementary Fig. 4A). Next, we repeated the GSA to determine the biological processes that are altered upon HE of TF genes by comparing the cumulative TPM values of genes in the 16 KEGG gene sets between samples with TF gene HE and Rest (Supplementary Fig. 4B). Compared with Rest, GSA identified significant associations of only *FOXA1* and *MYC* HE with the SRP gene set (Fig. 4B). It is worth noting that for both primary PCa and CRPCa, GSA provided consistent results only for sample stratification by *FOXA1* expression and to a minor extent by *MYC* expression. (Figs. 2C and 4B).

**Fig. 4:**
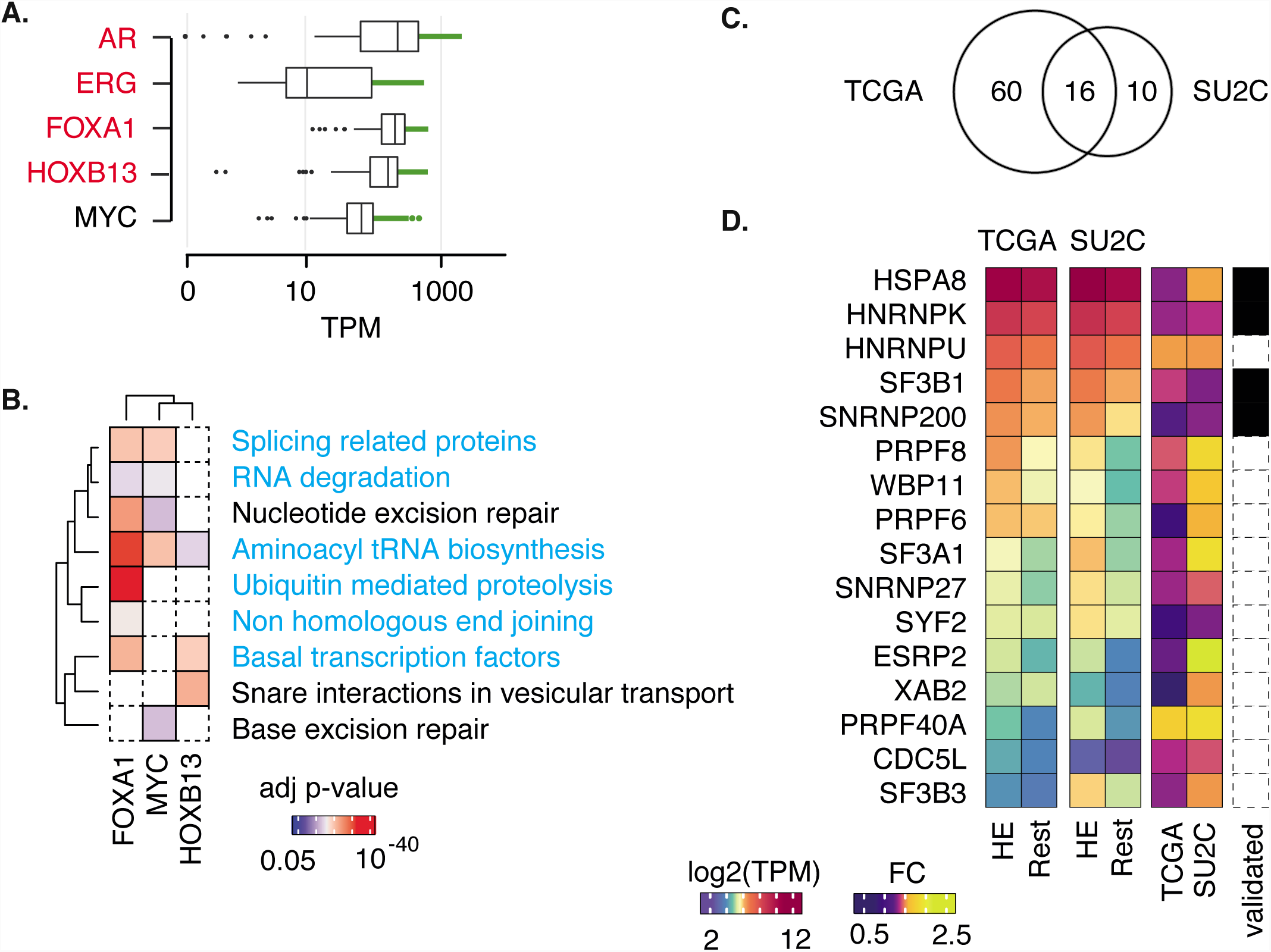
FOXA1 expression is independently associated with SRP expression in metastatic PCa. (A) Boxplot distributions of normalized gene expression levels (transcript per million reads, TPMs) for each of the five TFs in metastatic CRPCa samples (SU2C dataset). Green lines and dots refer to sample with TF expression greater than the 75th percentile of its distribution across samples. PCa-related TFs are reported in red. (B) Statistically significant results from the gene set analysis (GSA) for 16 biological KEGG processes related to the high expression (HE) of each TF. The altered processes are hierarchically clustered on the basis of their statistical significance (i.e. adjusted p-value). Non-statistically significant processes are depicted in white. The side panel reports the corresponding KEGG pathway category. Gene sets that were significantly altered by the five TFs in *FOXA1* HE PCa samples are reported in cyan. (C) Venn diagram showing the overlap between 76 and 26 significantly altered SRPs in PC and CRPCa samples, respectively, upon *FOXA1* HE. (D) Heatmap showing TPM values of altered SRPs in both primary PCa (TGCA) and CRPCa (SU2C) samples upon *FOXA1* HE and relative fold changes (FC). SRPs validated *in vitro* in PCa cell lines are reported in black. Missing values are depicted by white dashed blocks.

To determine whether FOXA1 regulates expression of the same SRP genes in both primary PCa and CRPCa, we compared significantly altered SRP genes associated with HE *FOXA1* in the TGCA and SU2C datasets (Fig. 4C, see Methods). We identified 16 SRP genes associated with HE *FOXA1* in both primary PCa and CRPCa (Fig. 4D). These data demonstrate the persistence of the FOXA1-SRP association in gene expression in treatment-relapsed CRPCa, which includes a subset of SRP genes (*HSPA8, HNRNPK, SF3B1* and *SNRNP200*) that both phenocopy FOXA1 and are regulated by FOXA1 *in vitro* (Figs. 2 and 3).

## Discussion

FOXA1 is a member of a group of forkhead box domain-containing transcription factors which interacts with chromatin and binds DNA, causing nucleosome rearrangement and stabilising an open chromatin conformation ^32^. FOXA1 has been described as a pioneer factor ^28^ for other transcription factors allowing the re-defining of cistromes ^7,30,33,34^, including the AR cistrome during prostate tumourigenesis ^7,35^. *FOXA1* is one of the most commonly mutated genes in PCa ^36,37^ and disease susceptibility loci fall within the FOXA1 cistrome ^10^, thereby highlighting its dysregulation in PCa.

Little is known of the mechanisms underpinning the transcriptional regulation of genes encoding RNA-binding and other proteins involved in AS ^11,16^, which we term as SRPs. To date, only MYC has been shown to transcriptionally drive expression of SRPs ^20-22^, thereby presenting AS as a cancer therapeutic vulnerability for *MYC*-driven tumours ^18,19^. Here, we show for the first time that FOXA1, but not AR or ERG, regulates a subset of SRP genes that phenocopy the *FOXA1* dependency of PCa cells, thereby expanding the AR-independent FOXA1 gene regulatory repertoire ^33,35^. AR-independent FOXA1-driven transcriptional programmes have been shown to occur via genomic interactions with other transcriptional regulators such as MYBL2 and CREB1 in CRPCa ^33^, GATA-3 and the Estrogen Receptor (ER) in breast cancer ^38^, and PPARγ in bladder cancer ^39^. Additionally, interactions between ER and glucocorticoid receptor (GR) and FOXA1 appear to be dynamic ^40^, hence our AR-independent observations may be context-dependent in PCa.

Considering the overlapping cistromes for FOXA1 and the ETS-family TF ERG ^30^, we were surprised that depletion of *ERG* and *FOXA1* had opposing effects on SRP gene expression. We concluded that this was due to an ERG-driven increase in *FOXA1* expression by the *ERG* siRNA. Overexpression of oncogenic ERG typically occurs as a result of a recurrent gene fusion between the AR-regulated *TMPRSS* gene and *ERG* in ∼50% PCa cases ^41^. Hence, our findings may highlight differing mechanisms of FOXA1-mediated SRP gene regulation in *ERG* fusion-positive and -negative PCa. Since loss of the tumour suppressor *PTEN* appears to be required for widespread co-operative AR and ERG-driven transcriptomic changes ^8^, we may have failed to identify ERG-driven SRP gene expression changes in PTEN-proficient VCaP cells. However, our data in PTEN-deficient LNCaP cells suggest that AR does not regulate SRP gene expression in the absence of PTEN.

A recent genome-wide screen of PCa cell dependencies ^42^ identified the heterogeneous nuclear ribonuclear protein (hnRNP) family of SRPs, including the FOXA1-regulated SRP genes *HNRNPA1* and *HNRNPK*, which have been previously implicated in PCa ^43-46^ and other malignancies^47^. The SRP hnRNPA1 has been shown to regulate expression of the CRPCa-and FOXA1-associated AR splice variant AR-V7 ^43,44,46,48,49^. Interestingly, we show that FOXA1 can control expression of *SF3B1*, which is the most-commonly altered SRP gene in PCa ^3^ and haematological malignancies ^50^. Lethality can be induced in cancers with mutations affecting *SF3B1* and other SRP genes by therapeutic targeting the SF3b complex of the spliceosome^51^, although the impact in cancers with up-regulated (but wild-type) *SF3B1* and other SRP genes is unknown. We also identify the SRP *HSPA8*, which encodes a heat shock protein (HSP) scaffold in the core spliceosome complex ^52^, as a FOXA1-regulated gene that exhibited a high dependency for PCa cells. Therapeutic targeting the HSP family member HSP90 has been shown to harbour anti-tumour activity and also modulate AS in CRPCa ^53,54^.

The role of FOXA1 in advanced PCa is still contradictory, with reports of AR-dependent ^55^ and -independent functions ^33,35^. On one hand, FOXA1 protein expression is up-regulated in primary PCa ^34,56,57^, metastases and CRPCa ^56^, and is associated with metastasis ^58^, disease recurrence ^56,57,59^, and survival ^34^. However, on the other hand, FOXA1 has also been described as an inhibitor of metastasis ^35^ and neuroendocrine differentiation ^60^, which is associated with a poor prognosis. We were unable to identify a difference in the levels of *FOXA1* expression amongst the FOXA1 HE tumours in the TCGA and SU2C datasets, however, we did observe a persistent association between *FOXA1* and SRP genes in CRPCa.

Our data demonstrate, for the first time in both primary PCa and CRPCa, that FOXA1 is associated with SRP gene expression, the dysregulation of which confers a poor patient prognosis. In a subset of FOXA1 binding and actively-transcribed SRP genes that phenocopy the *FOXA1* dependency of PCa cells, we confirm FOXA1-regulated SRP gene expression in PCa cell lines. Hence, we speculate that in both primary PCa and CRPCa, targeting SRPs may represent a therapeutic vulnerability for FOXA1-overexpressing PCa in an analogous way to *MYC*-driven cancers. This would need to be tested in future studies by therapeutic targeting of SRPs or upstream signaling cascades.

## Methods

### Gene set analysis of the human prostate cancer transcriptome

RNA sequencing (RNA-Seq) data were downloaded from The Cancer Genome Atlas (TCGA) Data Matrix portal (Level 3, https://tcga-data.nci.nih.gov/tcga/dataAccessMatrix.htm) and from cBioPortal^61,62^ websites for 409 primary untreated and 118 metastatic (Stand Up to Cancer, SU2C ^31^) PCa samples, respectively. The number of transcripts per million reads (TPM) was measured starting from the scaled estimate expression values provided for 20,531 genes as previously described ^63^. For the SU2C dataset RPKM (Reads Per Kilobase of transcript per Million mapped reads) values were converted into TPM. For each TF, the distribution of expression levels across samples was measured. A TF was considered as highly expressed (HE) if its TPM value was greater or equal to the 75^th^ percentile of the distribution ^63^ (Supplementary Data 1).

A list of 16 manually curated gene sets representing Genetic Information Processing (2.1-2.4) from the Kyoto Encyclopedia of Genes and Genomes (KEGG) were downloaded from MSigDb version 5 ^24^. A list of 66 additional RBPs available from the RNAcompete catalogue ^14^ were added to the KEGG spliceosome gene set (n=128). To further refine a list of genes encoding SRPs, a gene ontology (GO) analysis of biological processes was performed using the function *gprofiler* in the R ‘gProfileR’ ^64^ on the total set of 194 genes. A final gene set of 148 genes with GO terms related to splicing (SRPs) was retained for further analyses.

For each TF *t* and each gene set *i*, the cumulative TPM values of genes in *i* were compared between samples where the *t* was HE and the remaining ones (Rest) using a two-tailed Wilcoxon rank sum test (Supplementary Fig. 1A). The resulting p-values were corrected for multiple tests using the Bonferroni method. To control for false discoveries, Monte Carlo simulation was implemented as previously described ^65^. For 10,000 times, we randomly extracted 103 samples (corresponding to the number of samples with HE TF) and the cumulative expression of each gene set was compared with that of the remaining samples. Next, for each gene set, the empirical p-value is measured as the number of times the p-value is smaller than the observed one over the total number of iterations.

### Patient survival analysis

Clinical data were downloaded from the TCGA Data Matrix portal. Disease-free survival time was defined as the interval between the date of treatment and disease progression, as defined by biochemical or clinical recurrence, or until end of follow-up ^2^. The relationship with disease recurrence for genes within the top 6 gene sets *i* significantly associated with HE of all TFs was tested using a multivariable Cox proportional hazards (PH) model, and coefficients αα for each gene *j* were used to calculate a patient gene set score as following:

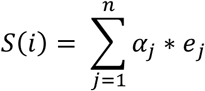

where e is the expression level of each gene.

For each gene set, patients were then stratified on the 75^th^ percentile of the score distribution, and a univariable Cox PH model was used to generate hazard ratios (HR) between patients with a high score as compared to those with a low score. For the SRP gene set scores, event-time distributions for the time to disease progression were compared using the log-rank test. The PH assumption and influential observations were met. All analyses were performed using the R ‘survival’ package ^66^.

### Linear regression modelling of the SRP gene set

SRP gene expression was fitted on the expression of the TFs using a linear regression model. The search for the best subsets of regressor was performed using a branch-and-bound algorithm ^67^ implemented in the *regsubsets* function in th R ‘leaps’ package. For models using a different number of variables (i.e. from 1 to 5 TFs) the best model in terms of determination coefficient R^2^ was reported. Relative importance of regressors in the linear regression model of five TFs was calculated using the function *calc.relimp* in the R ‘reclaimpo’ package ^68^. This function divides the determination coefficient R^2^ into the contribution of each regressor using the averaging over ordering method ^25^. The confidence intervals of the contributions of the regressors were measured using a bootstrap procedure implemented in the function *boot.relaimp*. For 1,000 iterations the full observation vectors were resampled and the regressor contributions were calculated.

### Selection of highly expressed SRPs

SRPs that were highly expressed between *FOXA1* HE (n=103 and 30 for TCGA and SU2C, respectively) and Rest (n=306 and 88 for TCGA and SU2C, respectively) were identified comparing the TPM distributions of the two groups with a one-tailed Kolmogorov-Smirnov (KS) test. The resulting p-values were corrected for multiple tests using the Bonferroni method. To control for size differences between the two cohorts, a Monte Carlo procedure was implemented. For 10,000 times, *FOXA1* HE and Rest samples were randomly selected and, for each SRP, the TPM distributions were compared using a one-tailed KS test. Next, for each SRP gene, the empirical p-value was measured as the proportion of tests with p-value smaller than the corresponding observed one over the total number of iterations. Differentially expressed genes (DEG) between *FOXA1* HE and Rest samples were detected using the R package ‘DESeq2’ and ‘EdgeR’ for the TCGA dataset, for which raw sequencing counts were available. Briefly, read counts of 20,531 genes of each sample were used as input for DESeq2 and EdgeR. Genes with read count equal to zero across all samples were removed. For the TCGA dataset, a total of 76 SRP genes with a KS Bonferroni-corrected p-value and empirical p-value less than 0.05, a false discovery rate (FDR) ≤ 0.1 and an absolute log_2_ Fold-Change (FC) ≥ 0.2 measured by DESeq2 or EdgeR were considered as having an altered expression in *FOXA1* HE samples as compared to Rest. For the SU2C dataset, a total of 26 SRP genes with a KS Bonferroni-corrected p-value and empirical p-value less than 0.05 and an absolute log_2_ (FC) ≥ 0.2 were considered as altered (Supplementary Data 2).

### Selection of TFs implicated in SRP gene expression

Enrichment analysis of TF binding sites within SRP genes was performed using chromatin immunoprecipitation sequencing (ChIP-Seq) data from the ReMap database and annotation tool ^69^. Genomic coordinates of SRP genes were extended by 5,000 bp and used as input for the ReMap enrichment tools (http://tagc.univ-mrs.fr/remap/index.php?page=annotation). Binding sites of TFs that significantly overlapped (minimum 10%) with the input regions were collected. For all five TFs, binding sites in regions of active transcription as defined by H3K27ac and H3K4me3 epigenetic modification markers were further selected. In particular, H3K27ac ChIP-Seq replicated narrow peaks from prostate epithelial cells were collected from the ENCODE Data Matrix (https://www.encodeproject.org/files/ENCFF655JIF/) and mapped on hg19 using The University of California, Santa Cruz (UCSC) liftover software (http://genome-euro.ucsc.edu/cgi-bin/hgLiftOver). H3K4me4 ChIP-Seq narrow peaks from LNCaP cells were retrieved from the UCSC hg19 database (http://hgdownload.soe.ucsc.edu/goldenPath/hg19). TF binding site regions were retained if overlapping with at least 10% of their sequence with H3K27ac and H3K4me3 peaks (Supplementary Data 2).

### Cell dependency analysis

Gene dependency data for 17,098 genes across 4 PCa cell lines (DU145, PC3, LNCaP, VCaP) ^27^ were downloaded from Project Achilles data portal (https://portals.broadinstitute.org/achilles). DEMETER inferred z-scores representing gene knockdown effects were extracted for *FOXA1* and SRP genes.

### Cell lines, antibodies, plasmids, oligonucleotides

DU145 (HTB-81, ATCC), PC3 (CRL-1435, ATCC), LNCaP (CRL-1740, ATCC), and VCaP (CRL-2876, ATCC) cells were obtained from American Type Culture Collection (ATCC) and identities confirmed by Short Tandem Repeat (STR) profiling (DDC Medical). pcDNA3.1 FOXA1 was provided by Jason Carroll (Cancer Research UK Cambridge Institute). The following antibodies were used: anti-FOXA1 (Abcam: ab23738), anti-actin (Sigma: A1978), anti-AR (BD Bioscience: 554225), anti-ERG (Santa Cruz: sc-271048), anti-mouse IgG HRP-linked (Dako: P044701-2), anti-rabbit IgG HRP-linked (Dako: P044801-2). Sequences used to generate siRNA duplexes are as previously described^70^ or commercially-designed (ON-TARGETPlus, Dharmacon Horizon Discovery) and are listed in Supplementary Table 3. Sequences used to generate oligonucleotide primers for PCR were designed by entering the Ensembl (http://www.ensembl.org) Transcript ID representing the principal isoform for each gene into the University Probe Library (UPL) Assay Design Centre (https://lifescience.roche.com/en_gb/brands/universal-probe-library.html#assay-design-center). Sequences were checked using the National Center for Biotechnology Information (NCBI) Primer-BLAST tool (https://www.ncbi.nlm.nih.gov/tools/primer-blast) prior to synthesis (Integrated DNA Technologies). Primer sequences are listed in Supplementary Table 8.

### Cell Culture, DNA and RNA transfections

Cells were incubated at 37°C, 5% CO_2_ in a humidified incubator. Cells were maintained at sub-confluency in RPMI-1640 medium (21875-034, Gibco) (DU145, PC3 and LNCaP) or DMEM (41966-029, Gibco) (VCaP) containing 2mM L-glutamine, supplemented with 10% foetal calf serum (FCS) (Gibco), 100 units/ml penicillin and 100 µg/ml streptomycin (15140-122, Gibco) and regularly tested for the presence of mycoplasma. Transfections with plasmid DNA and siRNA duplexes were carried out as detailed in the figure legends using ViaFect (E4981, Promega) and RNAiMax (13778-075, Thermo Fisher Scientific), respectively, according to manufacturers’ instructions.

### Sodium Dodecyl Sulphate PolyAcrylamide Gel Electrophoresis (SDS-PAGE) and Western blotting

Whole cell lysate protein samples were obtained by lysis of cells in RIPA (Radio-Immunoprecipitation Assay) buffer for 30 minutes at 4°C followed by lysate clearing by centrifugation. Protein concentration was calculated using the bicinchoninic acid (BCA) assay (10678484, Thermo Fisher Scientific) method and samples adjusted to equal concentrations of total protein. Samples were denatured in a 2-Mercapto-ethanol-based SDS sample buffer. Proteins were then separated by SDS-PAGE, transferred onto PVDF (polyvinylidene difluoride) membrane (000000003010040001, Sigma) using the wet transfer method, blocked in 5% milk in TBST (Tris-Buffered Saline and Polysorbate 20) and then placed in primary antibodies diluted in 5% BSA (Bovine Serum Albumin) in TBST over-night at 4°C. Membranes were washed and incubated with relevant HRP-conjugated secondary antibodies for 1 hour at room temperature. For signal detection, membranes were washed and incubated for 3 minutes each in Luminata Crescendo Western HRP substrate (10776189, Thermo Fisher Scientific) before bands were visualised on a Chemidoc system (Amersham Imager 600, Amersham). Antibody concentrations were as follows: anti-FOXA1 (1:1000), anti-actin (1:100,000), anti-AR (1:1000), anti-ERG (1:1000); HRP-linked secondaries (1:5000). Where indicated, densitometric assessments of protein bands were performed using Image Studio Lite Ver 5.2 (LI-COR), and signal intensities used to calculate relative normalised FC in protein expression.

### RNA extraction and quantitative reverse transcription polymerase chain reaction (qRT-PCR)

Total RNA was isolated from cells by direct lysis in TRIzol Reagent (15596026, Thermo Fisher Scientific) according to the manufacturer’s instructions and contaminating genomic DNA removed using DNase I (AMPD1-1KT, Sigma). Reverse transcription (RT) to cDNA was achieved using the High-Capacity cDNA Reverse Transcription Kit (4368813, Thermo Fisher Scientific). qRT-PCR was performed on the 7500 Fast Real-Time PCR machine (Applied Biosystems, 4351106) using triplicate cDNA templates with the FastStart Universal Probe Master with ROX (4913949001, Roche) and UPL set (04683633001, Roche) according to the manufacturer’s instructions. Only primers within a 10% efficiency range from 90-100% were included (Supplementary Table 8). Reaction conditions were as follows: 20 s at 50 °C, 10 min at 95 °C, and 40 cycles of 15 s at 95 °C and 1 min at 60 °C. Relative gene expression was determined by the 2^-ΔΔCT^ method using the geometric mean of two validated endogenous control genes (*ACTB* and *B2M*) to ensure the reliability and reproducibility of observed effects. Data shown are from three independent biological experimental replicates with two technical replicates.

### Cell viability assays

Cell viability assays were performed using (3-(4,5-Dimethylthiazol-2-yl)-2,5-Diphenyltetrazolium Bromide) (MTT) (L11939.06, Alfa Aesar) according to the manufacturer’s instructions. Briefly, 4000– 10 000 cells were seeded into each well of a 96-well plate and grown to ∼20–30% confluence prior to transfection with siRNA. After 72 h, MTT was added to each well to a final concentration of 0.67 mg/ml and incubated at 37°C, 5% CO_2_ in a humidified incubator for 2 h. Subsequently, MTT reagent was removed, 100µl dimethyl sulfoxide (DMSO) (10213810, Thermo Fisher Scientific) added to each well and agitated at room temperature for 15 mins. Absorbance was measured at 560nm and 630nm (SpectraMax Plus384 Absorbance Microplate Reader, Molecular Devices), and normalised by subtracting the 630nm value from the 560nm value, and percentage viability calculated as follows: Treatment absorbance ÷ DMSO control absorbance ×100. All siRNA data were normalized to a non-silencing control. Results shown are the means ± SEM of at least three independent experiments with at least 3 technical replicates.

### Statistical analysis for in vitro data

Graphical data shown represent the means ± standard error of the mean (SEM) of independent experiments. The one-tailed independent sample t-test was employed to identify differences in means between groups with p-value < 0.05 taken to indicate statistical significance.

## Acknowledgements

The following reagents were generously gifted: VCaP cells (from Y-J. Lu, Barts Cancer Institute, UK), and pcDNA3-FOXA1 (from J. Carroll, Cancer Research UK Cambridge Institute, UK). This work was funded by a joint Royal College of Surgeons of England/Cancer Research UK Clinician Scientist Fellowship in Surgery (C19198/A15339 to PR), The Urology Foundation and John Black Charitable Foundation (to PR), the Barts Charity, the Orchid Charity (to JDK and NL) and by the Italian Association for Cancer Research (AIRC MFAG 20566 to MC and IG 20240 to SO).

## Author Contributions

JGF, MC and PR designed research. JGF, RA, MDG, CA, AL, and MC performed research. JDK, NRL and SO contributed new reagents or analytic tools. JGF, MDG, CA, MC, and PR analysed data. JGF, MC and PR wrote the paper.

## Materials & Correspondence

Correspondence and material requests should be addressed to either

