## Supplementary Information for "Dysregulation of splicing-related proteins in prostate cancer is controlled by FOXA1"

**This Word file includes:**

Supplementary Figs. 1 to 5

Supplementary Tables 1 to 8

Captions for Supplementary Data 1 and 2

**Other supplementary materials for this manuscript include the following:**

Datasets S1 to S2

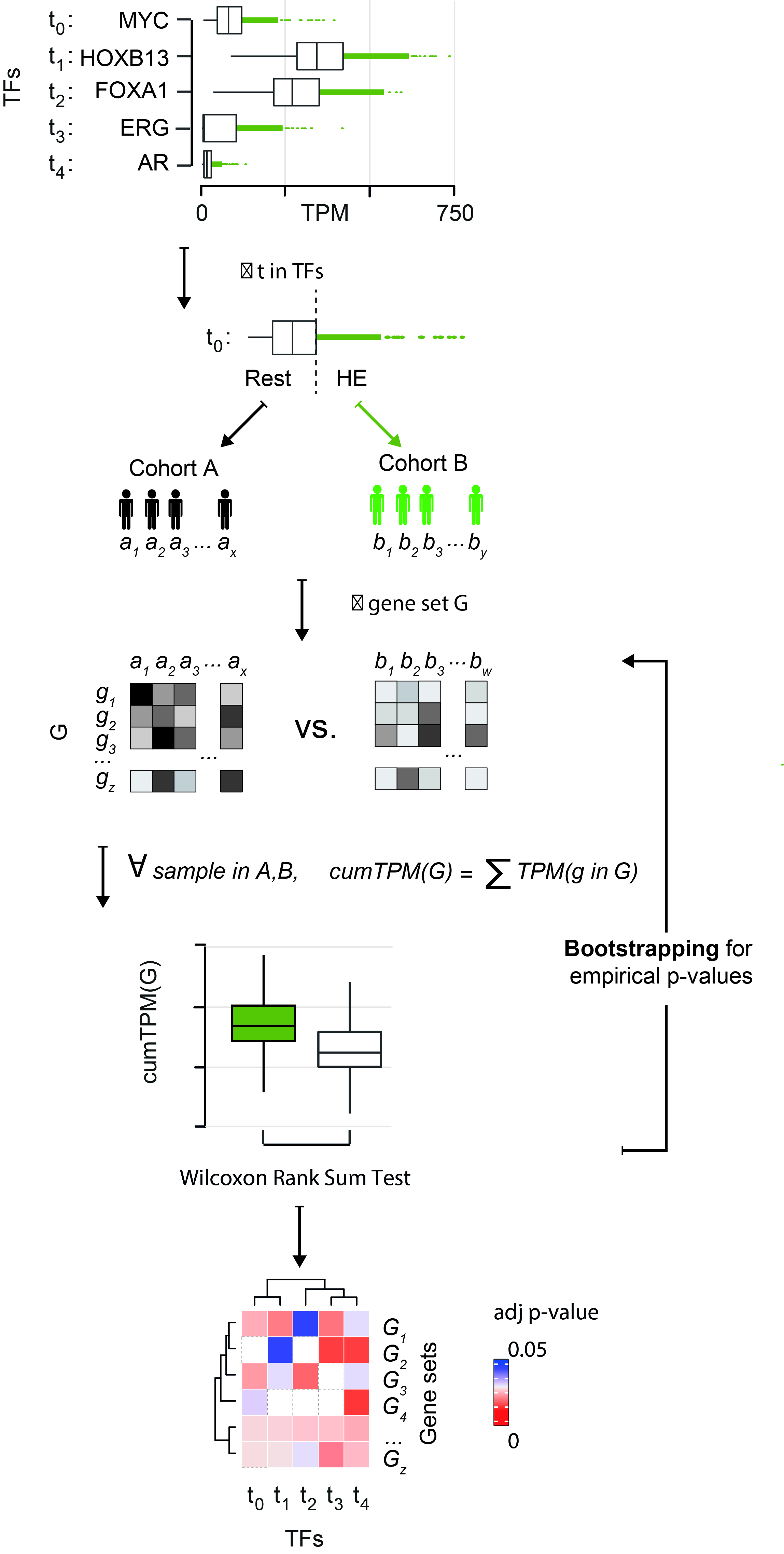

Supplementary Fig. 1. Overview of the gene set analysis (GSA) method. Samples were stratified according to the expression of each TF. In particular, highly expressed samples (HE) were defined as those having the expression of a TF higher than the 75th percentile of the TF expression distribution across all samples. For each TF stratification and each gene set, the distributions of cumulative expression levels of the genes in a gene set were compared between HE and the remaining ones (Rest) samples using Wilcoxon two-tailed Rank-Sum test. p-values were then corrected using Bonferroni adjustment. A bootstrapping procedure was implemented to estimate the empirical significance of the obtained p-values.

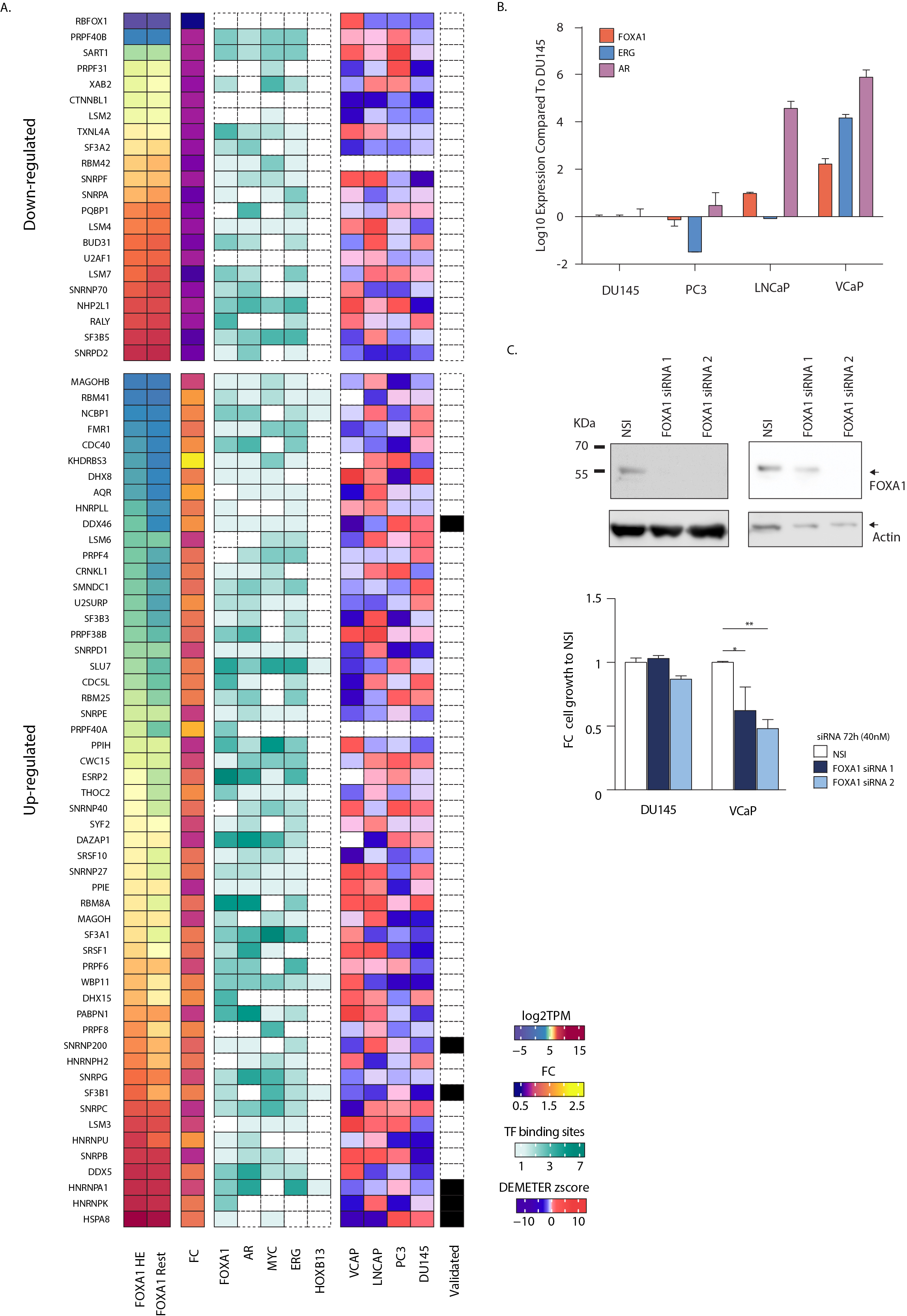

Supplementary Fig. 2. (A) Significantly dysregulated SRPs in *FOXA1* HE vs Rest patient samples are displayed, ranked by TPM from *FOXA1* HE samples. ReMap analysis was used, as described in the Methods to calculate enrichment values which are presented for five TFs: FOXA1, AR, MYC, ERG and HOXB13 for each SRP. DETEMER z-scores were retrieved from Project Achilles for each SRP. (B) qRT-PCR was performed on cDNAs from different PCa cells lines, and levels of *FOXA1*, *ERG*, *AR*, and *HOXB13* transcript expression were normalized to a geometric mean of *ACTB* and *B2M* levels to calculate Log_10_ relative normalized fold change (FC) in expression compared to DU145 cells. Data from at least three independent experiments were used to calculate the means ± SEM (C) DU145 and VCaP cells were transfected with two siRNA duplex sequences to *FOXA1* (FOXA1 si1 or si2), or non-silencing (NSI) control to final concentration of 40 nM. After 72 hours: (Upper panel) protein samples were collected and western blotting performed to confirm FOXA1 knockdown and (Lower panel) a MTT assay was performed to assess cell growth. Data from at least three independent experiments were used to calculate the means ± SEM.

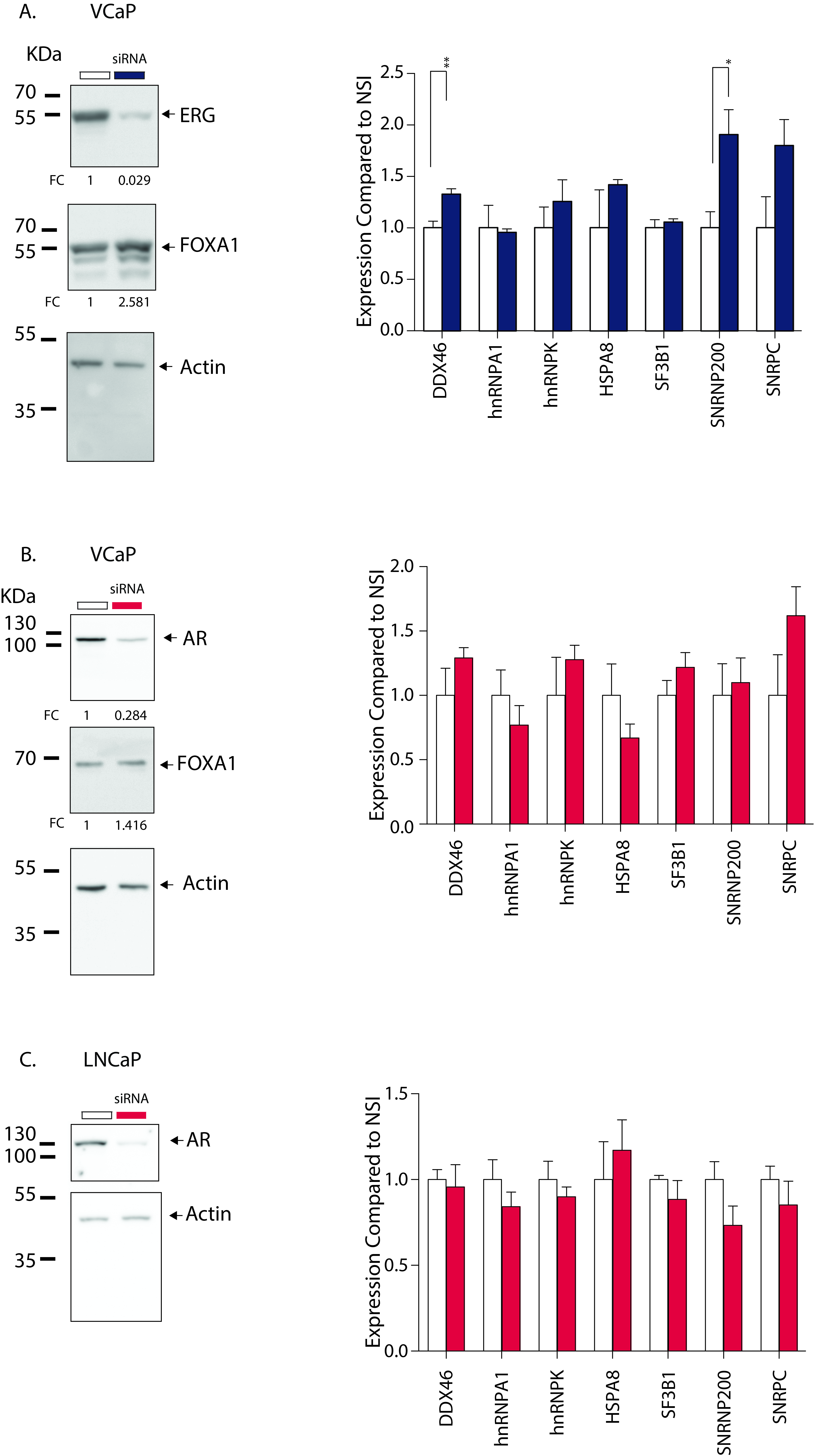

Supplementary Fig. S3. (A) VCaP cells were transfected with a pool of 4 siRNA duplex sequences to ERG (siERG), or non-silencing (NSI) control to final concentration of 40 nM. (B) VCaP and (C) LNCaP cells were transfected with a single siRNA duplex sequences to *AR* (siAR), or non-silencing (NSI) control to final concentration of 40 nM. After 72 h, total cell lysates were harvested and subjected to western blotting with antibodies to AR and actin. Western blotting images shown are representative of three independent experiments, from which densitometric band quantitation was performed to calculate the mean relative normalized fold change (FC) in protein expression (shown in brackets). (A-C, right panels) qRT-PCR was performed on cDNAs and levels of SRP transcript expression were normalized to a geometric mean of *ACTB* and *B2M* levels to calculate relative normalised FC in expression compared to NSI. Data from at least three independent experiments were used to calculate the means ± SEM. Unpaired two-tailed T-test was used to compared groups: *p-value <0.05, **p-value <0.01, ***p-value <0.001, ****p-value <0.0001

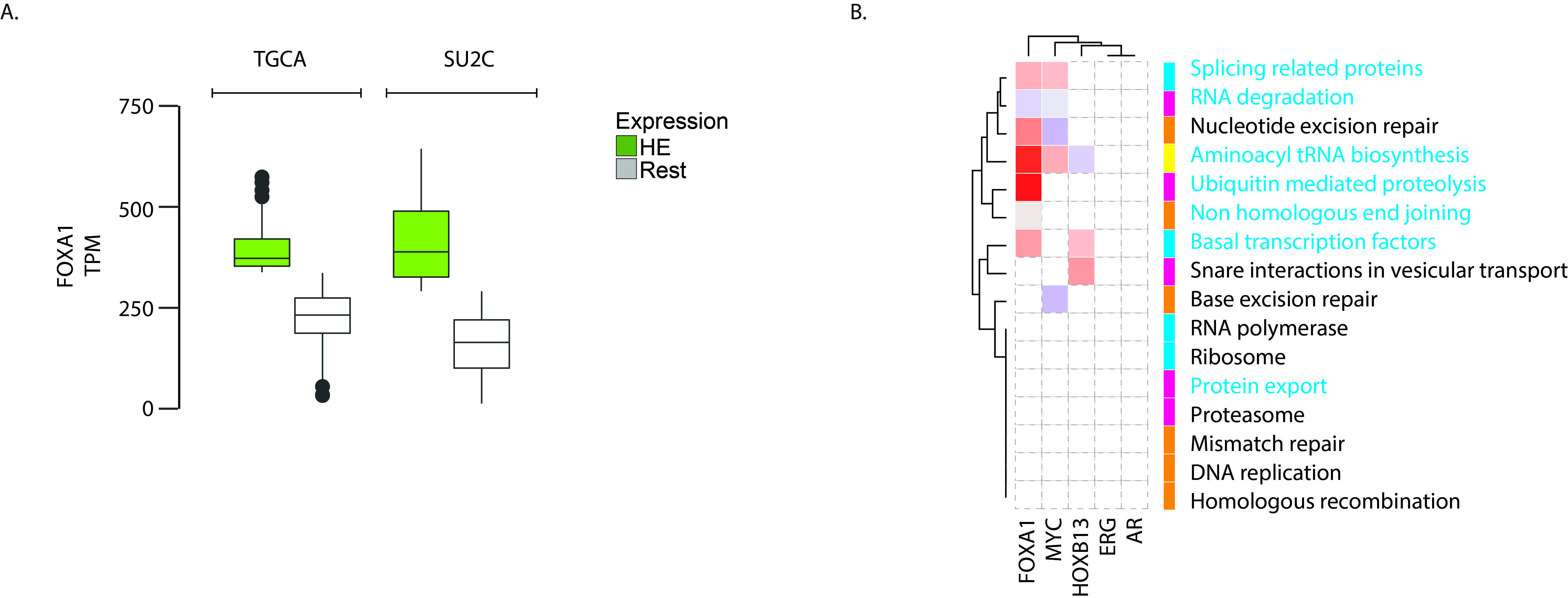

Supplementary Fig 4. (A) Comparison of *FOXA1* expression as TPM for *FOXA1* HE vs Rest stratification in the TGCA and SU2C datasets. (B) Results from the gene set analysis (GSA) for 16 biological KEGG processes related to the HE of each TF. The altered processes are hierarchically clustered on the basis of their statistical significance (i.e. adjusted p-value). Non-statistically significant processes are depicted in white. The side panel reports the corresponding KEGG pathway category.

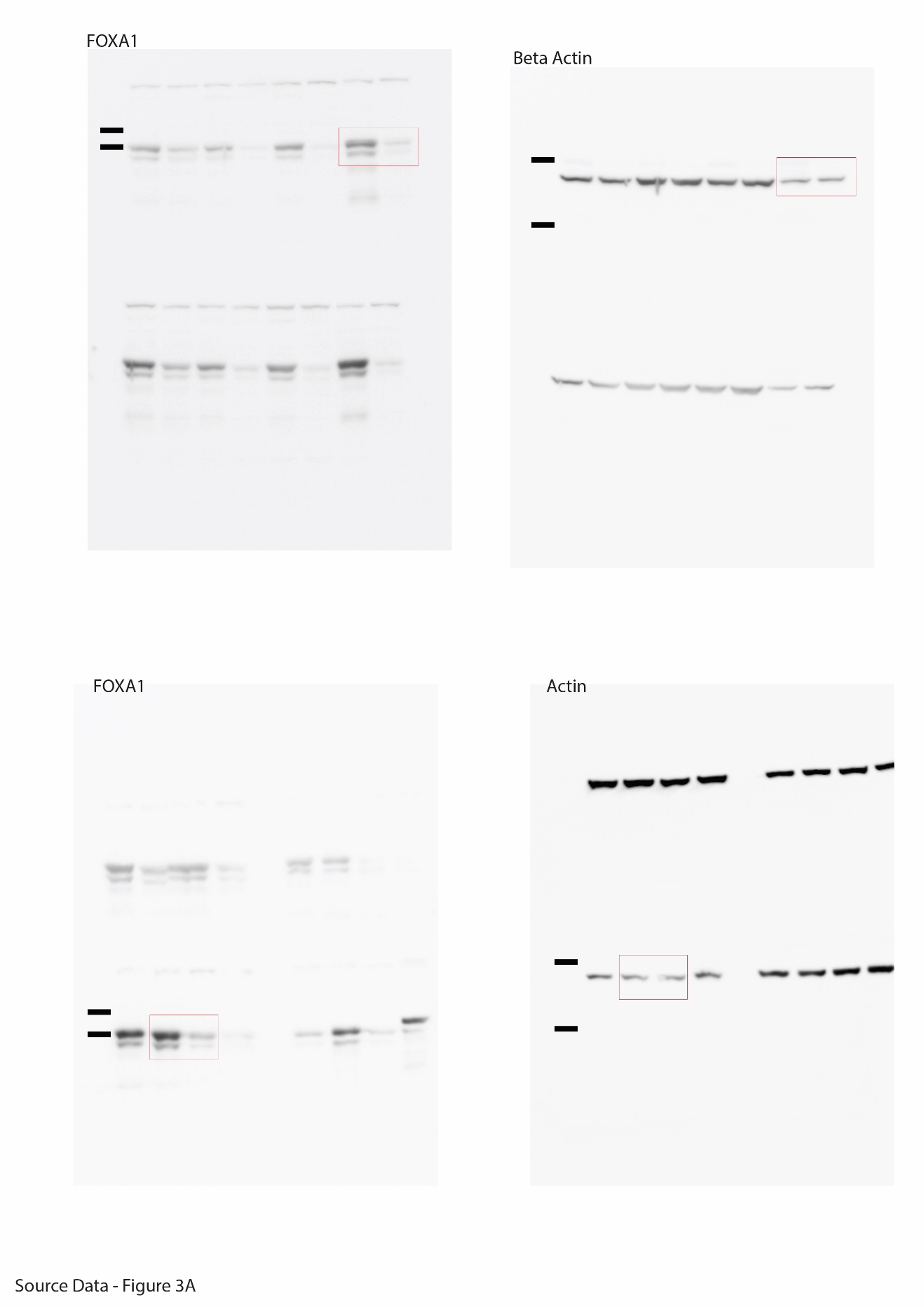

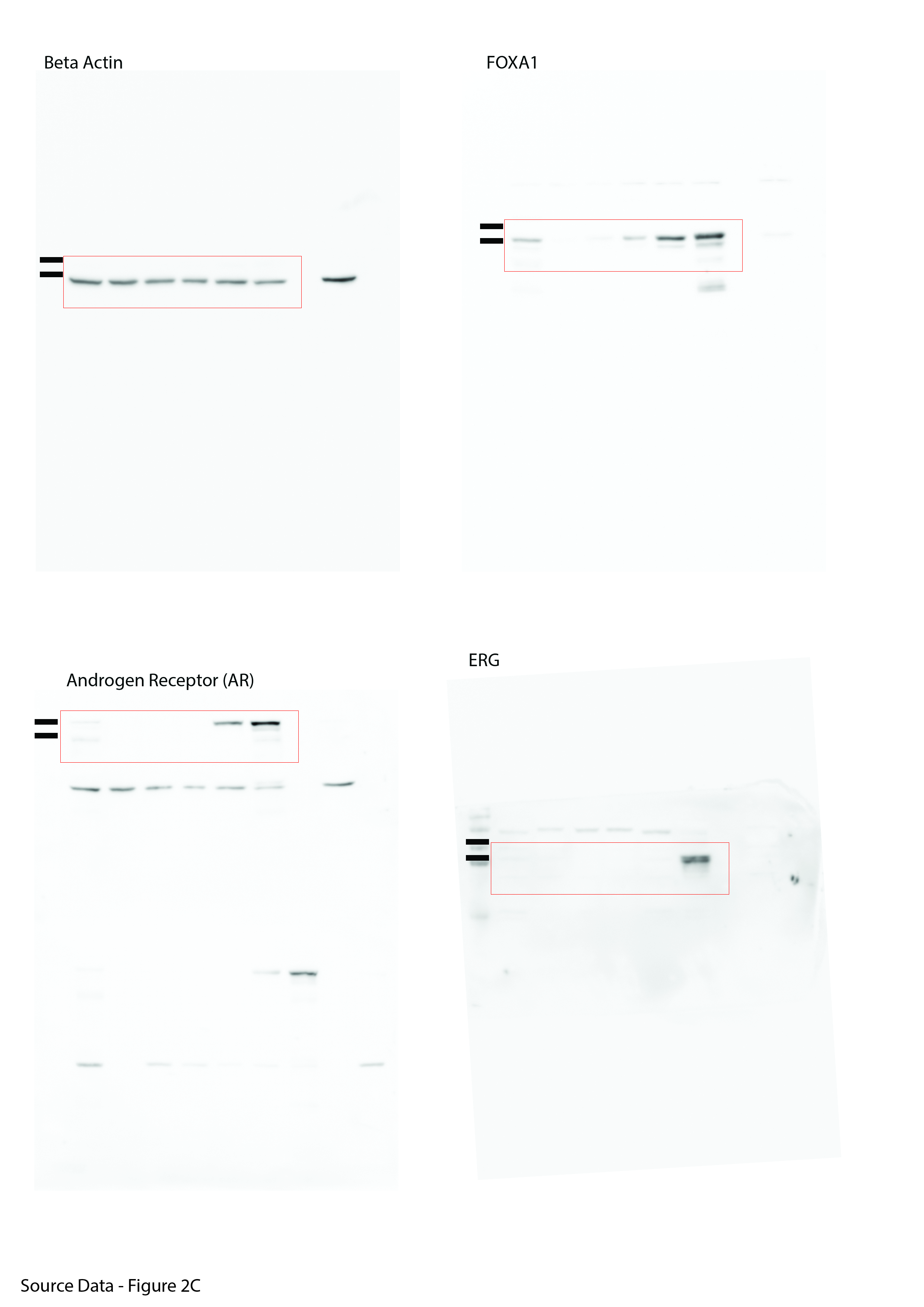

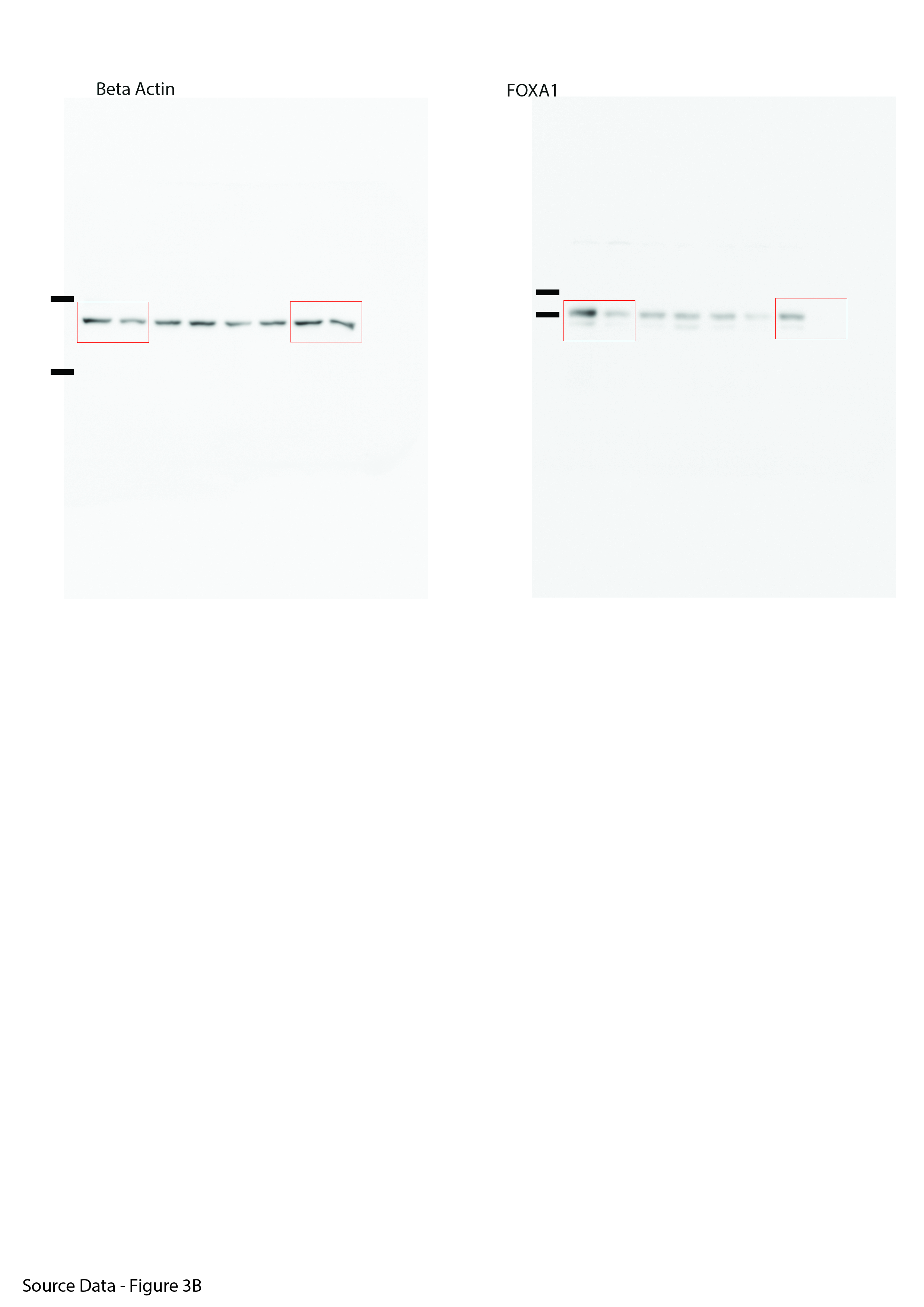

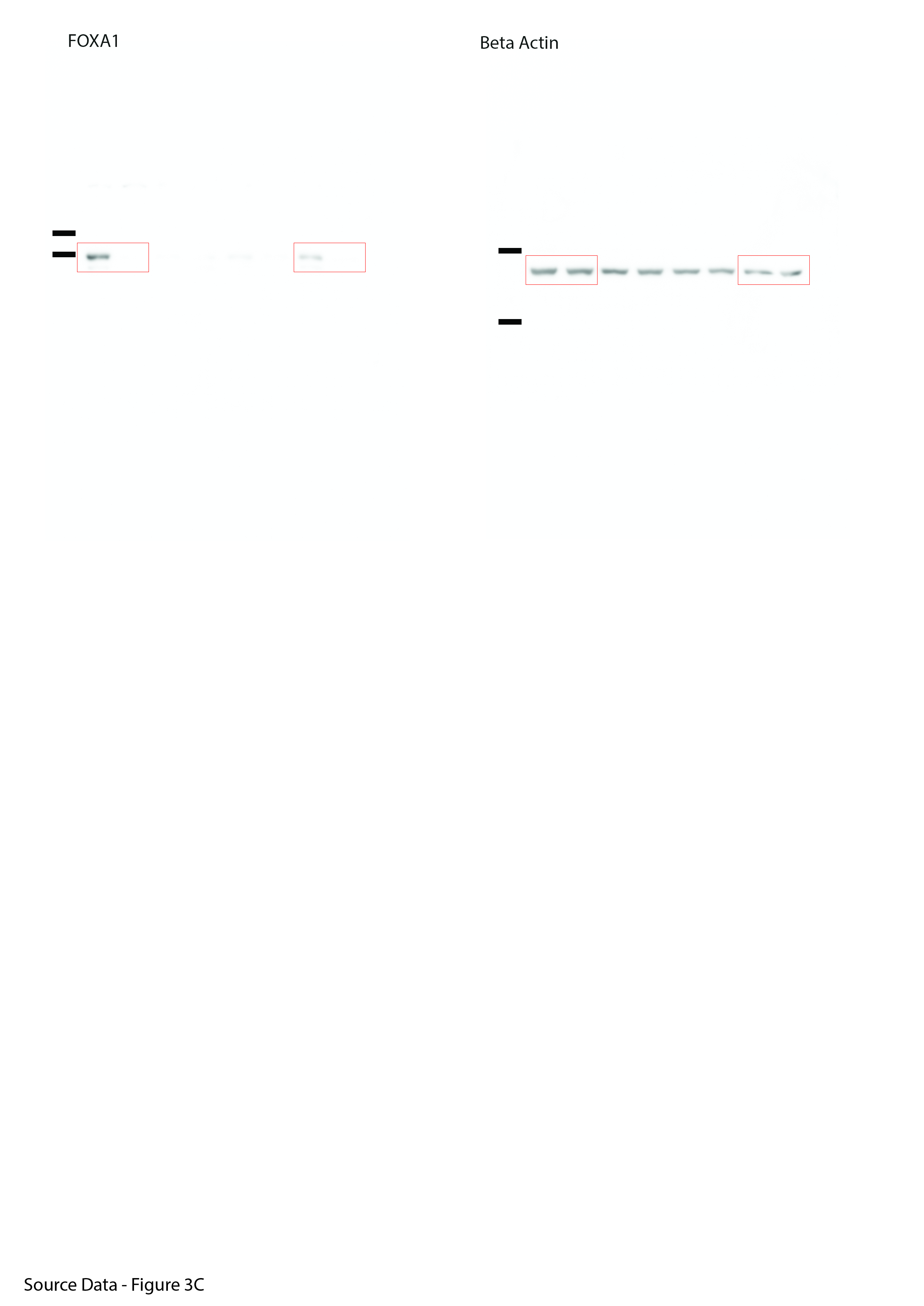

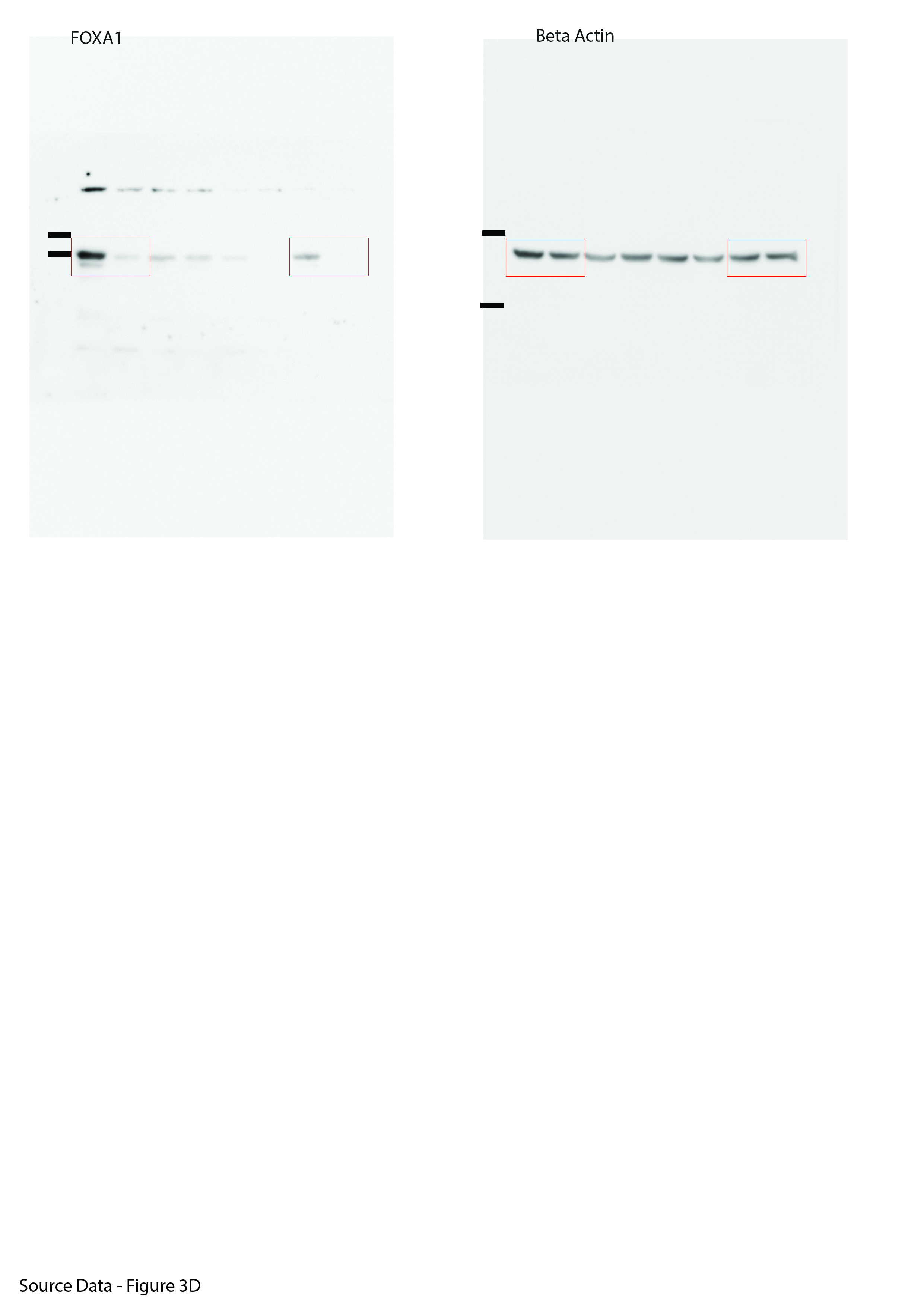

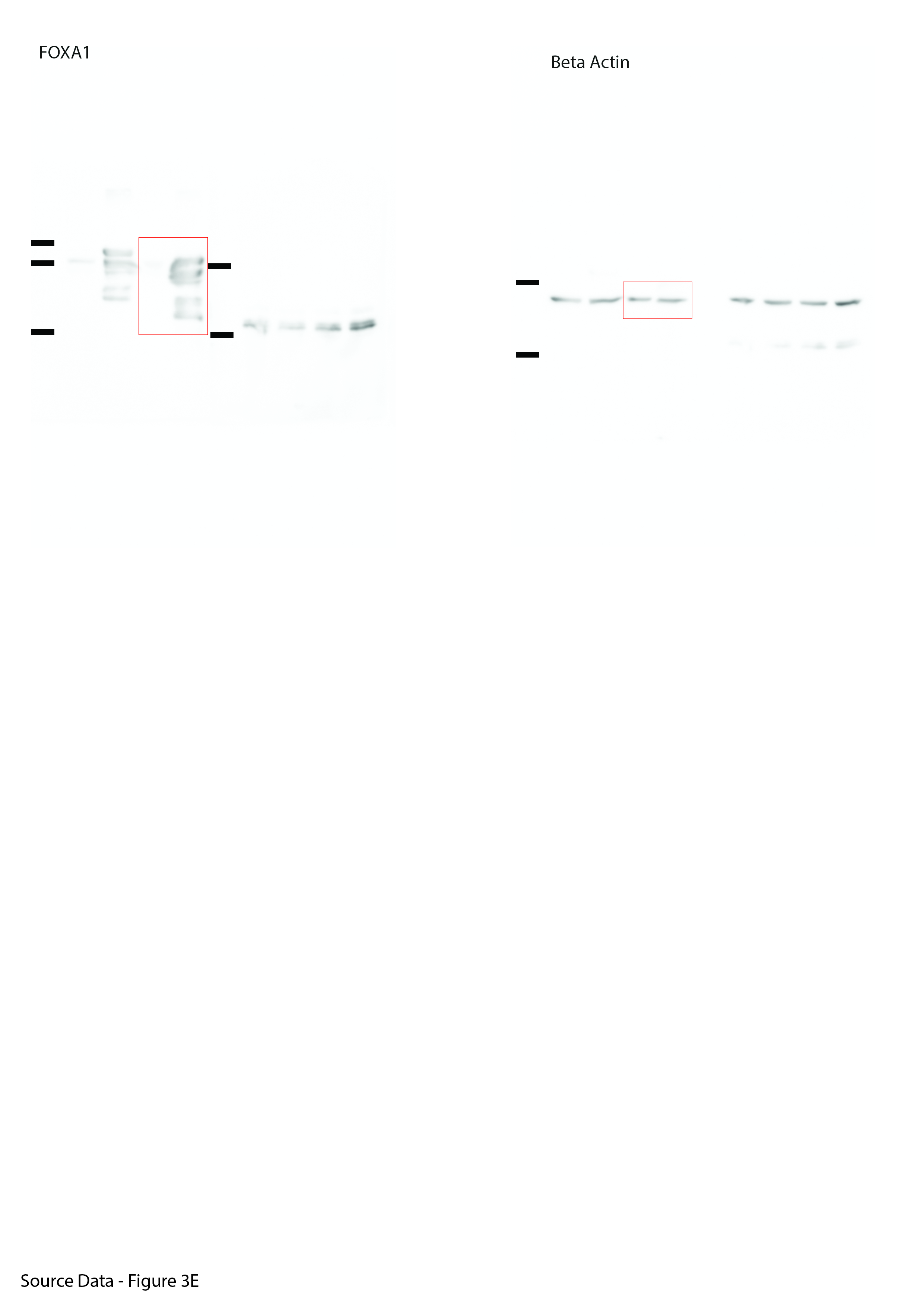

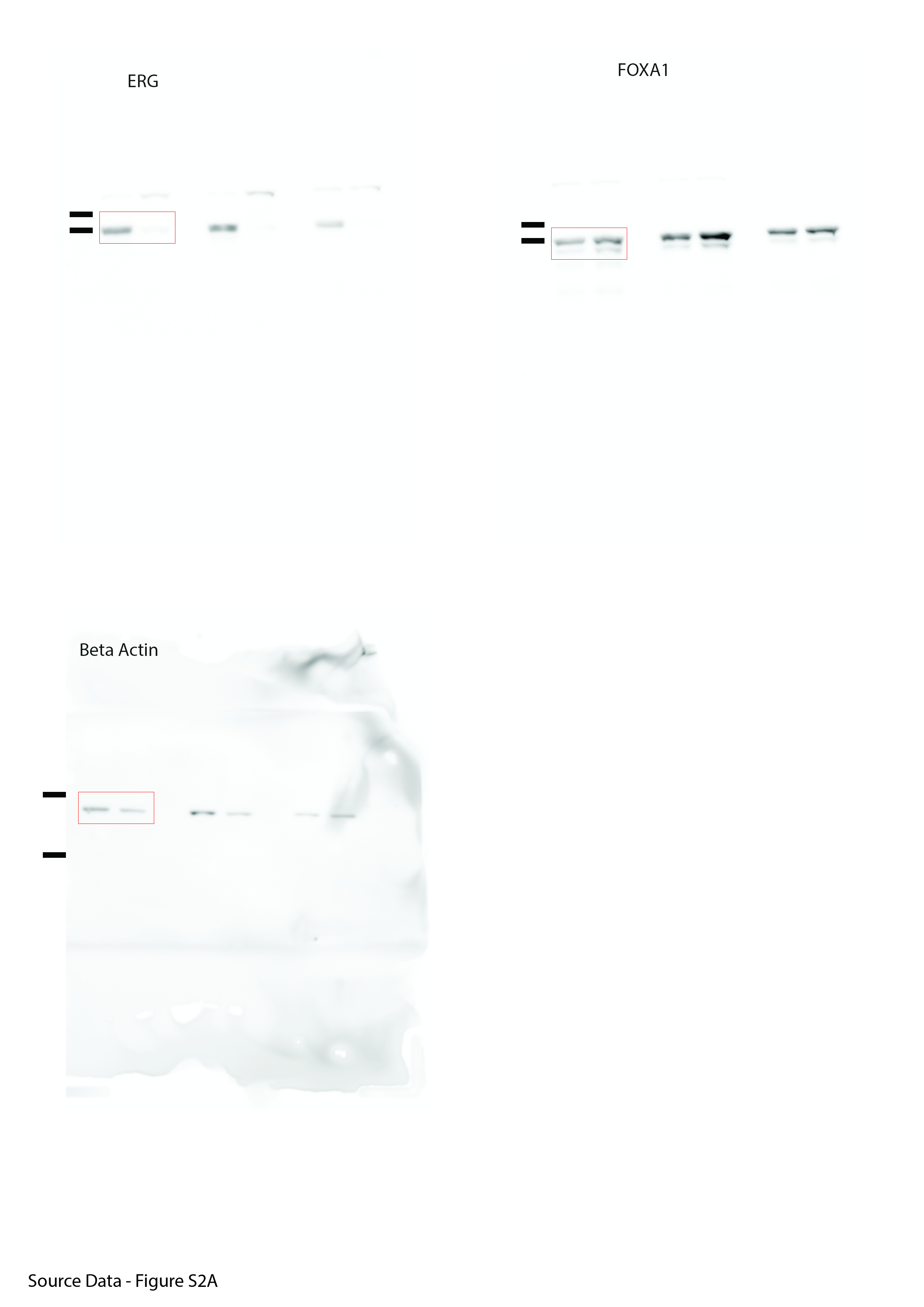

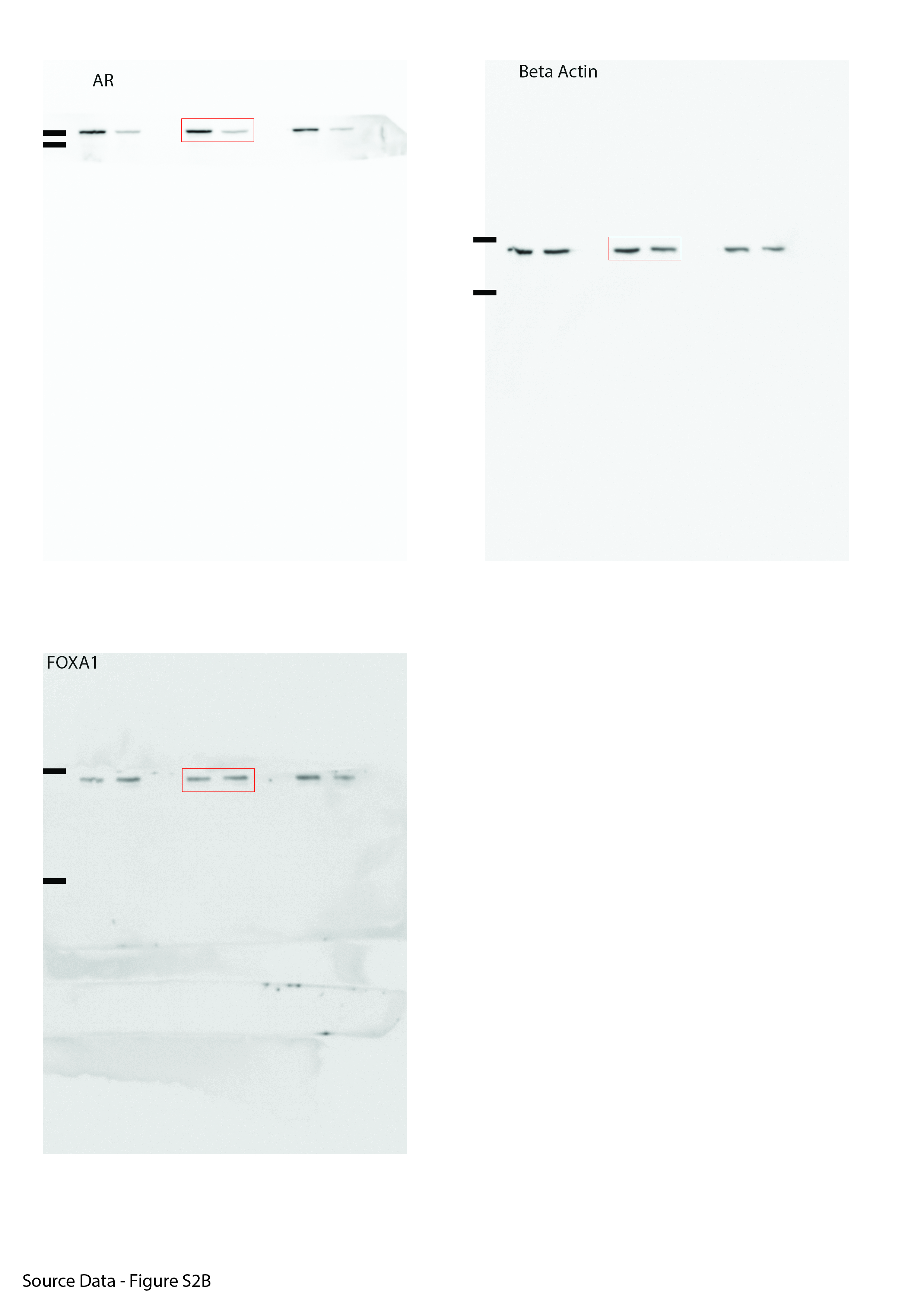

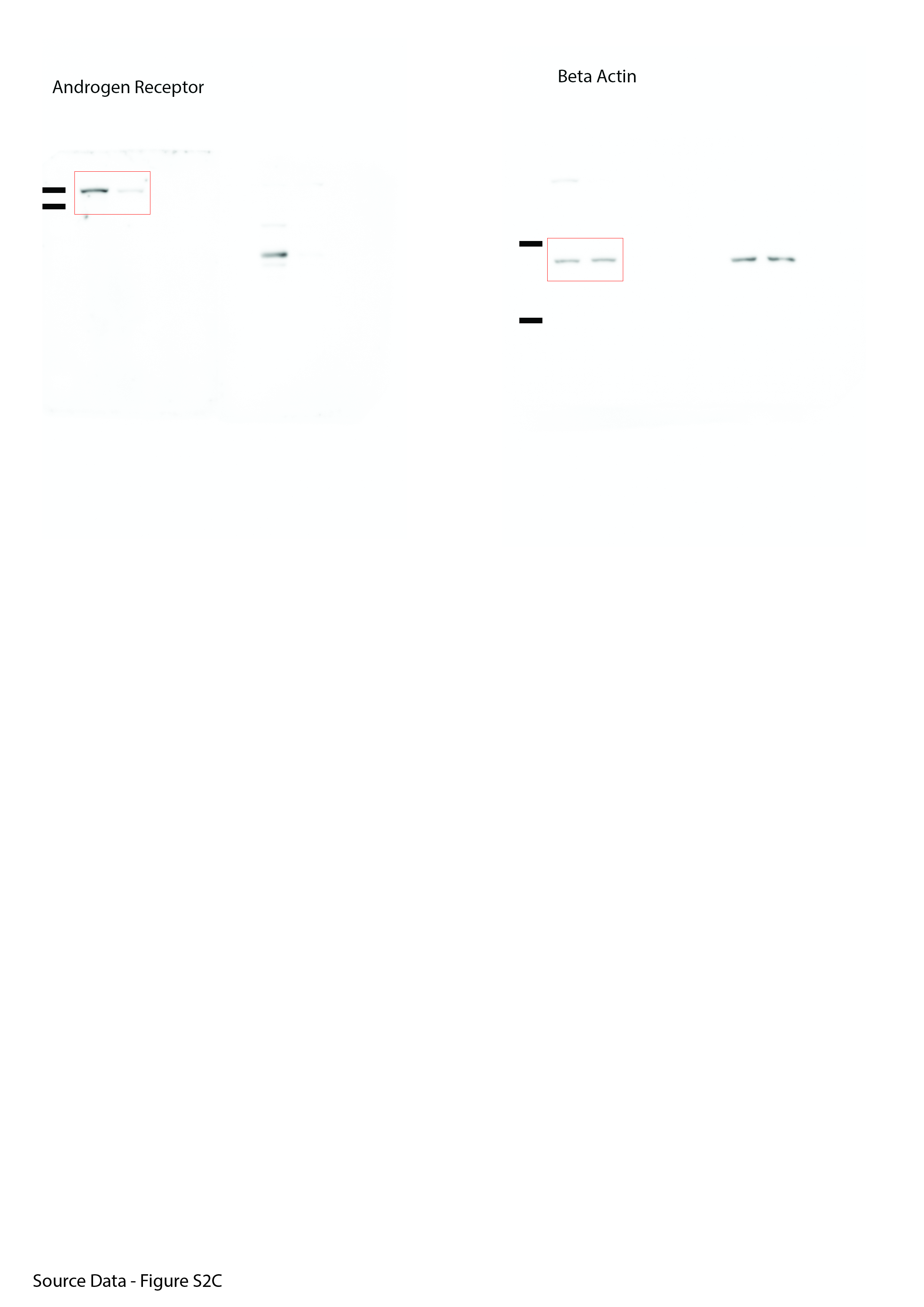

Supplementary Fig. S5. Full size western blot images with red boxes to indicate the cropped images used in the indicated figure panels as representative images

Supplementary Table 1. Results of Wilcoxon test results for the gene set analysis (GSA) for 16 biological KEGG processes following transcription factor stratification

| **condition** | **gene_set** | **statistic** | **p.value** | **bonf** | **p.emp** |
| --- | --- | --- | --- | --- | --- |
| MYC_overexpression | AMINOACYL_TRNA_BIOSYNTHESIS | 14613 | 1.94E-23 | 1.28E-21 | 1.00E-04 |
| MYC_overexpression | BASAL_TRANSCRIPTION_FACTORS | 13998 | 2.53E-19 | 1.49E-17 | 1.00E-04 |
| MYC_overexpression | BASE_EXCISION_REPAIR | 10924 | 4.24E-05 | 1.99E-03 | 1.95E-02 |
| MYC_overexpression | DNA_REPLICATION | 9407 | 7.04E-02 | 1.00E+00 | 2.70E-01 |
| MYC_overexpression | HOMOLOGOUS_RECOMBINATION | 5735 | 1.00E+00 | 1.00E+00 | 9.99E-01 |
| MYC_overexpression | MISMATCH_REPAIR | 9825 | 1.58E-02 | 5.28E-01 | 1.82E-01 |
| MYC_overexpression | NON_HOMOLOGOUS_END_JOINING | 14432 | 3.49E-22 | 2.23E-20 | 1.00E-04 |
| MYC_overexpression | NUCLEOTIDE_EXCISION_REPAIR | 11653 | 1.60E-07 | 8.63E-06 | 1.13E-02 |
| MYC_overexpression | PROTEASOME | 7597 | 9.28E-01 | 1.00E+00 | 9.94E-01 |
| MYC_overexpression | PROTEIN_EXPORT | 10911 | 4.63E-05 | 2.13E-03 | 3.12E-01 |
| MYC_overexpression | RIBOSOME | 8702 | 3.71E-01 | 1.00E+00 | 4.13E-02 |
| MYC_overexpression | RNA_DEGRADATION | 14056 | 1.08E-19 | 6.70E-18 | 1.00E-04 |
| MYC_overexpression | RNA_POLYMERASE | 8189 | 6.92E-01 | 1.00E+00 | 7.12E-01 |
| MYC_overexpression | SNARE_INTERACTIONS_IN_VESICULAR_TRANSPORT | 9314 | 9.31E-02 | 1.00E+00 | 8.81E-01 |
| MYC_overexpression | SPLICING_RELATED_PROTEINS | 14010 | 2.13E-19 | 1.30E-17 | 1.00E-04 |
| MYC_overexpression | UBIQUITIN_MEDIATED_PROTEOLYSIS | 13552 | 1.33E-16 | 7.42E-15 | 2.00E-04 |
| HOXB13_overexpression | AMINOACYL_TRNA_BIOSYNTHESIS | 15139 | 2.68E-27 | 1.82E-25 | 1.00E-04 |
| HOXB13_overexpression | BASAL_TRANSCRIPTION_FACTORS | 14302 | 2.64E-21 | 1.64E-19 | 1.00E-04 |
| HOXB13_overexpression | BASE_EXCISION_REPAIR | 11817 | 3.78E-08 | 1.82E-06 | 1.00E-04 |
| HOXB13_overexpression | DNA_REPLICATION | 9695 | 2.62E-02 | 5.51E-01 | 2.15E-02 |
| HOXB13_overexpression | HOMOLOGOUS_RECOMBINATION | 6308 | 1.00E+00 | 1.00E+00 | 9.07E-01 |
| HOXB13_overexpression | MISMATCH_REPAIR | 10075 | 5.31E-03 | 1.33E-01 | 1.01E-02 |
| HOXB13_overexpression | NON_HOMOLOGOUS_END_JOINING | 14290 | 3.18E-21 | 1.94E-19 | 1.00E-04 |
| HOXB13_overexpression | NUCLEOTIDE_EXCISION_REPAIR | 13060 | 7.28E-14 | 3.79E-12 | 1.00E-04 |
| HOXB13_overexpression | PROTEASOME | 10656 | 2.36E-04 | 8.26E-03 | 1.00E-04 |
| HOXB13_overexpression | PROTEIN_EXPORT | 13674 | 2.52E-17 | 1.38E-15 | 1.00E-04 |
| HOXB13_overexpression | RIBOSOME | 5998 | 1.00E+00 | 1.00E+00 | 1.00E+00 |
| HOXB13_overexpression | RNA_DEGRADATION | 14202 | 1.22E-20 | 7.06E-19 | 1.00E-04 |
| HOXB13_overexpression | RNA_POLYMERASE | 9649 | 3.11E-02 | 6.00E-01 | 2.00E-03 |
| HOXB13_overexpression | SNARE_INTERACTIONS_IN_VESICULAR_TRANSPORT | 12534 | 3.10E-11 | 1.55E-09 | 1.00E-04 |
| HOXB13_overexpression | SPLICING_RELATED_PROTEINS | 13695 | 1.88E-17 | 1.06E-15 | 1.00E-04 |
| HOXB13_overexpression | UBIQUITIN_MEDIATED_PROTEOLYSIS | 15088 | 6.54E-27 | 4.32E-25 | 1.00E-04 |
| FOXA1_overexpression | AMINOACYL_TRNA_BIOSYNTHESIS | 15499 | 4.03E-30 | 2.58E-28 | 1.00E-04 |
| FOXA1_overexpression | BASAL_TRANSCRIPTION_FACTORS | 15498 | 4.11E-30 | 2.59E-28 | 1.00E-04 |
| FOXA1_overexpression | BASE_EXCISION_REPAIR | 11506 | 5.48E-07 | 1.86E-05 | 6.00E-04 |
| FOXA1_overexpression | DNA_REPLICATION | 10107 | 4.57E-03 | 9.02E-02 | 1.60E-02 |
| FOXA1_overexpression | HOMOLOGOUS_RECOMBINATION | 6062 | 1.00E+00 | 1.00E+00 | 9.87E-01 |
| FOXA1_overexpression | MISMATCH_REPAIR | 10637 | 2.65E-04 | 6.88E-03 | 3.20E-03 |
| FOXA1_overexpression | NON_HOMOLOGOUS_END_JOINING | 14975 | 4.62E-26 | 2.77E-24 | 1.00E-04 |
| FOXA1_overexpression | NUCLEOTIDE_EXCISION_REPAIR | 13360 | 1.67E-15 | 9.04E-14 | 1.00E-04 |
| FOXA1_overexpression | PROTEASOME | 9712 | 2.46E-02 | 3.20E-01 | 3.62E-02 |
| FOXA1_overexpression | PROTEIN_EXPORT | 13339 | 2.20E-15 | 1.16E-13 | 1.00E-04 |
| FOXA1_overexpression | RIBOSOME | 5883 | 1.00E+00 | 1.00E+00 | 1.00E+00 |
| FOXA1_overexpression | RNA_DEGRADATION | 14239 | 6.94E-21 | 3.89E-19 | 1.00E-04 |
| FOXA1_overexpression | RNA_POLYMERASE | 8449 | 5.32E-01 | 1.00E+00 | 5.14E-01 |
| FOXA1_overexpression | SNARE_INTERACTIONS_IN_VESICULAR_TRANSPORT | 12544 | 2.78E-11 | 1.28E-09 | 1.00E-04 |
| FOXA1_overexpression | SPLICING_RELATED_PROTEINS | 14807 | 7.94E-25 | 4.61E-23 | 1.00E-04 |
| FOXA1_overexpression | UBIQUITIN_MEDIATED_PROTEOLYSIS | 15744 | 3.99E-32 | 2.72E-30 | 1.00E-04 |
| ERG_overexpression | AMINOACYL_TRNA_BIOSYNTHESIS | 14643 | 1.19E-23 | 7.49E-22 | 1.00E-04 |
| ERG_overexpression | BASAL_TRANSCRIPTION_FACTORS | 14659 | 9.16E-24 | 5.86E-22 | 1.00E-04 |
| ERG_overexpression | BASE_EXCISION_REPAIR | 10816 | 8.65E-05 | 3.72E-03 | 3.02E-02 |
| ERG_overexpression | DNA_REPLICATION | 8112 | 7.34E-01 | 1.00E+00 | 9.80E-01 |
| ERG_overexpression | HOMOLOGOUS_RECOMBINATION | 5429 | 1.00E+00 | 1.00E+00 | 1.00E+00 |
| ERG_overexpression | MISMATCH_REPAIR | 9070 | 1.77E-01 | 1.00E+00 | 7.55E-01 |
| ERG_overexpression | NON_HOMOLOGOUS_END_JOINING | 14907 | 1.47E-25 | 9.73E-24 | 1.00E-04 |
| ERG_overexpression | NUCLEOTIDE_EXCISION_REPAIR | 11603 | 2.45E-07 | 1.25E-05 | 1.94E-02 |
| ERG_overexpression | PROTEASOME | 8900 | 2.57E-01 | 1.00E+00 | 4.96E-01 |
| ERG_overexpression | PROTEIN_EXPORT | 12567 | 2.16E-11 | 1.17E-09 | 1.00E-04 |
| ERG_overexpression | RIBOSOME | 8884 | 2.66E-01 | 1.00E+00 | 5.80E-03 |
| ERG_overexpression | RNA_DEGRADATION | 14081 | 7.46E-20 | 4.48E-18 | 1.00E-04 |
| ERG_overexpression | RNA_POLYMERASE | 7936 | 8.19E-01 | 1.00E+00 | 8.70E-01 |
| ERG_overexpression | SNARE_INTERACTIONS_IN_VESICULAR_TRANSPORT | 10258 | 2.18E-03 | 7.40E-02 | 3.14E-01 |
| ERG_overexpression | SPLICING_RELATED_PROTEINS | 13976 | 3.49E-19 | 2.03E-17 | 1.00E-04 |
| ERG_overexpression | UBIQUITIN_MEDIATED_PROTEOLYSIS | 13798 | 4.47E-18 | 2.50E-16 | 1.00E-04 |
| AR_overexpression | AMINOACYL_TRNA_BIOSYNTHESIS | 15842 | 6.04E-33 | 3.98E-31 | 1.00E-04 |
| AR_overexpression | BASAL_TRANSCRIPTION_FACTORS | 15492 | 4.59E-30 | 2.76E-28 | 1.00E-04 |
| AR_overexpression | BASE_EXCISION_REPAIR | 11240 | 4.45E-06 | 1.42E-04 | 2.00E-03 |
| AR_overexpression | DNA_REPLICATION | 10529 | 5.00E-04 | 1.00E-02 | 7.00E-04 |
| AR_overexpression | HOMOLOGOUS_RECOMBINATION | 5405 | 1.00E+00 | 1.00E+00 | 1.00E+00 |
| AR_overexpression | MISMATCH_REPAIR | 11193 | 6.32E-06 | 1.90E-04 | 2.00E-04 |
| AR_overexpression | NON_HOMOLOGOUS_END_JOINING | 15658 | 2.05E-31 | 1.31E-29 | 1.00E-04 |
| AR_overexpression | NUCLEOTIDE_EXCISION_REPAIR | 13787 | 5.22E-18 | 2.82E-16 | 1.00E-04 |
| AR_overexpression | PROTEASOME | 8414 | 5.54E-01 | 1.00E+00 | 8.45E-01 |
| AR_overexpression | PROTEIN_EXPORT | 12976 | 2.01E-13 | 9.84E-12 | 1.00E-04 |
| AR_overexpression | RIBOSOME | 5527 | 1.00E+00 | 1.00E+00 | 1.00E+00 |
| AR_overexpression | RNA_DEGRADATION | 14433 | 3.44E-22 | 1.99E-20 | 1.00E-04 |
| AR_overexpression | RNA_POLYMERASE | 7901 | 8.33E-01 | 1.00E+00 | 8.83E-01 |
| AR_overexpression | SNARE_INTERACTIONS_IN_VESICULAR_TRANSPORT | 12610 | 1.35E-11 | 5.94E-10 | 1.00E-04 |
| AR_overexpression | SPLICING_RELATED_PROTEINS | 14290 | 3.18E-21 | 1.78E-19 | 1.00E-04 |
| AR_overexpression | UBIQUITIN_MEDIATED_PROTEOLYSIS | 15551 | 1.53E-30 | 9.51E-29 | 1.00E-04 |

Supplementary Table 2. Univariable Cox proportional hazards (PH) hazard ratios (HR) and standard error (SE) stratified by 75th percentile of patient gene set score distributions for KEGG processes

| **Gene set** | **SE** | **HR** | **95% CI** |
| --- | --- | --- | --- |
| SPLICING_RELATED | 0.28 | 25.54 | 14.65-44.51 |
| UBIQUITIN_MEDIATED_PROTEOLYSIS | 0.25 | 14.66 | 8.91-24.14 |
| AMINOACYL_TRNA_BIOSYNTHESIS | 0.21 | 4.35 | 2.86-6.61 |
| RNA_DEGRADATION | 0.21 | 4.23 | 2.8-6.4 |
| BASAL_TRANSCRIPTION_FACTORS | 0.21 | 3.91 | 2.58-5.91 |
| NON_HOMOLOGOUS_END_JOINING | 0.21 | 3.24 | 2.13-4.91 |

Supplementary Table 3. Sequences and sources of siRNA Oligonucleotides used in this study.

| **Gene** | **Species** | **Product Code** | **Target Sequence** | **Supplier/Source** |
| --- | --- | --- | --- | --- |
| Non-targeting Pool | N/A | D-001810-10-05 | UGGUUUACAUGUCGACUAA, UGGUUUACAUGUUGUGUGA, UGGUUUACAUGUUUUCUGA, UGGUUUACAUGUUUUCCUA | Dharmacon (ON-TARGETplus) |
| FOXA1 | Human | J-010319-05 | GCACUGCAAUACAUCGCCUU | Dharmacon (ON-TARGETplus) |
| FOXA1 | Human | J-010319-06 | CCUCGGAGCAGCAGCAUAA | Dharmacon (ON-TARGETplus) |
| FOXA1 | Human | J-010319-07 | GAACAGCUACUACGCAGAC | Dharmacon (ON-TARGETplus) |
| FOXA1 | Human | J-010319-08 | CCUAAACACUUCCUAGCUC | Dharmacon (ON-TARGETplus) |
| FOXA1 | Human | N/A | Sense: CCAUGAACACCUACAUGACCAUGAA55 Anti-sense: UUCAUGGUCAUGUAGGUGUUCAUGG55 | Zheng L. Int J Clin Exp Pathol 2015 Synthesised by Eurogentec |
| Negative Control (DS NC1) | N/A | 51-01-14-03 | Unknown - Not provided by supplier | Integrated DNA Technologies |
| AR | Human | hs.Ri.AR.13.2 | Unknown - Not provided by supplier | Integrated DNA Technologies |
| siGENOME Non-Targeting siRNA Pool #1 | N/A | D-001206-13-05 | UAGCGACUAAACACAUCAA, UAAGGCUAUGAAGAGAUAC, AUGUAUUGGCCUGUAUUAG, AUGAACGUGAAUUGCUCAA | Dharmacon (siGENOME) |
| ERG | Human | M-003886-01-0005 | GAUCCUACGCUAUGGAGUA, GUGAAUGGCUCAAGGAACU, GCGCUACGCCUACAAGUUC, GGACAGACUUCCAAGAUGA | Dharmacon (siGENOME) |

Supplementary Table 4. ddCT mean values and standard deviation for RT-qPCR results in Fig. 2F

|  | **DU145** | | | **PC3** | | |
| --- | --- | --- | --- | --- | --- | --- |
| **Gene** | **Mean** | **STDEV** | **N** | **Mean** | **STDEV** | **N** |
| DDX46 | 1 | 0.454469 | 4 | 0.872176 | 0.251291 | 4 |
| HNRNPA1 | 1 | 0.341065 | 4 | 0.83341 | 0.204908 | 4 |
| HNRNPK | 1 | 0.381877 | 4 | 0.87362 | 0.246806 | 4 |
| HSPA8 | 1 | 0.406734 | 4 | 1.371889 | 0.859596 | 4 |
| SF3B1 | 1 | 0.376958 | 4 | 1.170501 | 0.288426 | 4 |
| SNRNP200 | 1 | 0.43076 | 4 | 0.940742 | 0.247784 | 4 |
| SNRPC | 1 | 0.19608 | 4 | 1.740498 | 0.3994 | 4 |
|  | **LNCaP** | | | **VCaP** | | |
| **Gene** | **Mean** | **STDEV** | **N** | **Mean** | **STDEV** | **N** |
| DDX46 | 0.539473 | 0.183316 | 4 | 8.268299 | 2.320727 | 4 |
| HNRNPA1 | 1.303323 | 0.478218 | 4 | 7.567047 | 6.747047 | 4 |
| HNRNPK | 2.458542 | 0.785435 | 4 | 6.707606 | 2.364602 | 4 |
| HSPA8 | 5.490971 | 1.331967 | 4 | 22.0507 | 11.04457 | 4 |
| SF3B1 | 2.600457 | 1.063209 | 4 | 8.482004 | 1.877648 | 4 |
| SNRNP200 | 1.990728 | 0.758322 | 4 | 12.078 | 2.363563 | 4 |
| SNRPC | 2.471607 | 0.460155 | 4 | 5.266335 | 0.376788 | 4 |

Supplementary Table 5. Western Blot Quantification. Analysis of band intensity in western blot images used in manuscript figure panels where indicated

| **Figure** | **Repeat** | **Name** | **FOXA1** | **Beta Actin** | **Normalised FOXA1** | **FC To Control** |
| --- | --- | --- | --- | --- | --- | --- |
| 2C-D | 1 | DU145 | 164000 | 22100000 | 0.01 | 1.04 |
|  | 1 | PC3 | 1110000 | 17800000 | 0.06 | 8.72 |
|  | 1 | LNCaP | 23600000 | 20900000 | 1.13 | 157.92 |
|  | 1 | VCaP | 34700000 | 16200000 | 2.14 | 299.57 |
|  | 2 | DU145 | 199000 | 19200000 | 0.01 | 1.45 |
|  | 2 | PC3 | 952000 | 28200000 | 0.03 | 4.72 |
|  | 2 | LNCaP | 12000000 | 15300000 | 0.78 | 109.69 |
|  | 2 | VCaP | 41900000 | 5700000 | 7.35 | 1028.06 |
|  | 3 | DU145 | 89800 | 24500000 | 0.00 | 0.51 |
|  | 3 | PC3 | 1120000 | 14300000 | 0.08 | 10.95 |
|  | 3 | LNCaP | 6680000 | 20700000 | 0.32 | 45.13 |
|  | 3 | VCaP | 39000000 | 16200000 | 2.41 | 336.69 |
| 3A | 1 | NSI | 39500000 | 33200000 | 1.19 | 1.00 |
|  | 1 | siFOXA1 1 | 9520000 | 18500000 | 0.51 | 0.43 |
|  | 1 | NSI | 25100000 | 14200000 | 1.77 | 1.00 |
|  | 1 | siFOXA1 2 | 2360000 | 14200000 | 0.17 | 0.09 |
|  | 2 | NSI | 49300000 | 3890000 | 12.67 | 1.00 |
|  | 2 | siFOXA1 1 | 9710000 | 2810000 | 3.46 | 0.27 |
|  | 2 | NSI | 17600000 | 1560000 | 11.28 | 1.00 |
|  | 2 | siFOXA1 2 | 1450000 | 1070000 | 1.36 | 0.12 |
|  | 3 | NSI | 46100000 | 9910000 | 4.65 | 1.00 |
|  | 3 | siFOXA1 1 | 9030000 | 8230000 | 1.10 | 0.24 |
|  | 3 | NSI | 7120000 | 4940000 | 1.44 | 1.00 |
|  | 3 | siFOXA1 2 | 460000 | 16800000 | 0.03 | 0.02 |
|  | 4 | NSI | 19300000 | 33200000 | 0.58 | 1.00 |
|  | 4 | siFOXA1 1 | 10200000 | 38800000 | 0.26 | 0.45 |
|  | 4 | siFOXA1 2 | 3620000 | 31200000 | 0.12 | 0.20 |
|  | 5 | NSI | 9390000 | 38300000 | 0.25 | 1.00 |
|  | 5 | siFOXA1 1 | 1010000 | 22100000 | 0.05 | 0.19 |
|  | 5 | siFOXA1 2 | 205000 | 20500000 | 0.01 | 0.04 |
|  | 6 | NSI | 61700000 | 24000000 | 2.57 | 1.00 |
|  | 6 | siFOXA1 1 | 12000000 | 24200000 | 0.50 | 0.19 |
|  | 6 | siFOXA1 2 | 2270000 | 4060000 | 0.56 | 0.22 |
| 3B | 1 | NSI | 37000000 | 33700000 | 1.10 | 1.00 |
|  | 1 | siFOXA1 1 | 6340000 | 16000000 | 0.40 | 0.36 |
|  | 1 | NSI | 12100000 | 35600000 | 0.34 | 1.00 |
|  | 1 | siFOXA1 2 | 11400 | 24700000 | 0.00 | 0.00 |
|  | 2 | NSI | 49400000 | 8950000 | 5.52 | 1.00 |
|  | 2 | siFOXA1 1 | 5990000 | 18500000 | 0.32 | 0.06 |
|  | 2 | NSI | 41300000 | 35300000 | 1.17 | 1.00 |
|  | 2 | siFOXA1 2 | 5510000 | 21600000 | 0.26 | 0.22 |
|  | 3 | NSI | 61800000 | 34300000 | 1.80 | 1.00 |
|  | 3 | siFOXA1 1 | 9400000 | 10800000 | 0.87 | 0.48 |
|  | 3 | NSI | 28900000 | 6530000 | 4.43 | 1.00 |
|  | 3 | siFOXA1 2 | 9220000 | 24300000 | 0.38 | 0.09 |
| 3C | 1 | NSI | 47000000 | 23200000 | 2.03 | 1.00 |
|  | 1 | siFOXA1 1 | 6880000 | 7480000 | 0.92 | 0.45 |
|  | 1 | NSI | 8260000 | 12700000 | 0.65 | 1.00 |
|  | 1 | siFOXA1 2 | 47100 | 41900000 | 0.00 | 0.00 |
|  | 2 | NSI | 38100000 | 38800000 | 0.98 | 1.00 |
|  | 2 | siFOXA1 1 | 97400 | 38100000 | 0.00 | 0.00 |
|  | 2 | NSI | 11300000 | 21500000 | 0.53 | 1.00 |
|  | 2 | siFOXA1 2 | 206000 | 20600000 | 0.01 | 0.02 |
|  | 3 | NSI | 44200000 | 22700000 | 1.95 | 1.00 |
|  | 3 | siFOXA1 1 | 2510000 | 15200000 | 0.17 | 0.08 |
|  | 3 | NSI | 11000000 | 16900000 | 0.65 | 1.00 |
|  | 3 | siFOXA1 2 | 63800 | 40000000 | 0.00 | 0.00 |
| 3D | 1 | NSI | 45300000 | 46300000 | 0.98 | 1.00 |
|  | 1 | siFOXA1 1 | 1690000 | 29100000 | 0.06 | 0.06 |
|  | 1 | NSI | 5760000 | 27000000 | 0.21 | 1.00 |
|  | 1 | siFOXA1 2 | 32100 | 29200000 | 0.00 | 0.01 |
|  | 2 | NSI | 50400000 | 30400000 | 1.66 | 1.00 |
|  | 2 | siFOXA1 1 | 2570000 | 44600000 | 0.06 | 0.03 |
|  | 2 | NSI | 27900000 | 10900000 | 2.56 | 1.00 |
|  | 2 | siFOXA1 2 | 854000 | 42600000 | 0.02 | 0.01 |
|  | 3 | NSI | 78300000 | 63600000 | 1.23 | 1.00 |
|  | 3 | siFOXA1 1 | 2410000 | 34900000 | 0.07 | 0.06 |
|  | 3 | NSI | 27200000 | 25900000 | 1.05 | 1.00 |
|  | 3 | siFOXA1 2 | 973000 | 33800000 | 0.03 | 0.03 |
| 3E | 1 | VO | 4070000 | 16200000 | 0.25 | 1.00 |
|  | 1 | FOXA1 | 78700000 | 20000000 | 3.94 | 15.66 |
|  | 2 | VO | 172000 | 30400000 | 0.01 | 1.00 |
|  | 2 | FOXA1 | 2670000 | 25100000 | 0.11 | 18.80 |
|  | 3 | VO | 6030000 | 39900000 | 0.15 | 1.00 |
|  | 3 | FOXA1 | 34200000 | 40100000 | 0.85 | 5.64 |
| S3A | 1 | NSI | 38200000 | 16800000 | 2.27 | 1.00 |
|  | 1 | siERG | 114000 | 8840000 | 0.01 | 0.01 |
|  | 2 | NSI | 39600000 | 19800000 | 2.00 | 1.00 |
|  | 2 | siERG | 1030000 | 7430000 | 0.14 | 0.07 |
|  | 3 | NSI | 15700000 | 4440000 | 3.54 | 1.00 |
|  | 3 | siERG | 428000 | 10200000 | 0.04 | 0.01 |
|  | 1 | NSI | 11400000 | 16800000 | 0.68 | 1.00 |
|  | 1 | siERG | 17300000 | 8840000 | 1.96 | 2.88 |
|  | 2 | NSI | 20700000 | 19800000 | 1.05 | 1.00 |
|  | 2 | siERG | 34500000 | 7430000 | 4.64 | 4.44 |
|  | 3 | NSI | 19300000 | 4440000 | 4.35 | 1.00 |
|  | 3 | siERG | 18500000 | 10200000 | 1.81 | 0.42 |
| S3B | 1 | NSI | 29800000 | 22400000 | 1.33 | 1.00 |
|  | 1 | siAR | 6730000 | 22500000 | 0.30 | 0.22 |
|  | 2 | NSI | 29900000 | 21500000 | 1.39 | 1.00 |
|  | 2 | siAR | 6930000 | 13000000 | 0.53 | 0.38 |
|  | 3 | NSI | 22400000 | 12000000 | 1.87 | 1.00 |
|  | 3 | siAR | 3720000 | 8190000 | 0.45 | 0.24 |
|  | 1 | NSI | 2880000 | 22400000 | 0.13 | 1.00 |
|  | 1 | siAR | 5020000 | 22500000 | 0.22 | 1.74 |
|  | 2 | NSI | 3590000 | 21500000 | 0.17 | 1.00 |
|  | 2 | siAR | 4050000 | 13000000 | 0.31 | 1.87 |
|  | 3 | NSI | 6160000 | 12000000 | 0.51 | 1.00 |
|  | 3 | siAR | 2720000 | 8190000 | 0.33 | 0.65 |
| S3C | 1 | NSI | 36100000 | 14800000 | 2.44 | 1.00 |
|  | 1 | siAR | 5240000 | 16000000 | 0.33 | 0.13 |
|  | 2 | NSI | 18200000 | 11900000 | 1.53 | 1.00 |
|  | 2 | siAR | 2970000 | 37500000 | 0.08 | 0.05 |
|  | 3 | NSI | 1220000 | 33500000 | 0.04 | 1.00 |
|  | 3 | siAR | 104000 | 23600000 | 0.00 | 0.12 |

Supplementary Table 6. ddCT mean values and standard deviation for RT-qPCR results in Fig. 3 and Fig. S3.

|  |  |  | **NSI** | | | **FOXA1 si1** | | | **FOXA1 si2** | | |
| --- | --- | --- | --- | --- | --- | --- | --- | --- | --- | --- | --- |
| **Cell Line** | **Figure** | **Gene** | **Mean** | **STDEV** | **N** | **Mean** | **STDEV** | **N** | **Mean** | **STDEV** | **N** |
| **VCaP** | 3A | DDX46 | 1 | 0.555436 | 5 | 0.641476 | 0.16057 | 5 | 0.375303 | 0.074625 | 5 |
|  |  | hnRNPA1 | 1 | 0.127989 | 5 | 0.749626 | 0.114635 | 5 | 0.544261 | 0.090079 | 5 |
|  |  | hnRNPK | 1 | 0.193603 | 5 | 0.714125 | 0.409139 | 5 | 0.690151 | 0.294047 | 5 |
|  |  | HSPA8 | 1 | 0.271195 | 5 | 0.809083 | 0.123488 | 5 | 0.564628 | 0.086034 | 5 |
|  |  | SF3B1 | 1 | 0.131312 | 5 | 0.792954 | 0.030097 | 5 | 0.869144 | 0.1419 | 5 |
|  |  | SNRNP200 | 1 | 0.115093 | 5 | 0.707112 | 0.153935 | 5 | 0.401728 | 0.073807 | 5 |
|  |  | SNRNPC | 1 | 0.787158 | 5 | 0.566093 | 0.62686 | 5 | 0.398638 | 0.417662 | 5 |
| **LNCaP** | 3B | DDX46 | 1 | 0.145733 | 3 | 1.379537 | 0.325241 | 3 | 1.037243 | 0.22515 | 3 |
|  |  | hnRNPA1 | 1 | 0.181029 | 3 | 0.849672 | 0.116065 | 3 | 0.300245 | 0.049816 | 3 |
|  |  | hnRNPK | 1 | 0.168878 | 3 | 0.966179 | 0.096488 | 3 | 0.501575 | 0.116281 | 3 |
|  |  | HSPA8 | 1 | 0.212538 | 3 | 1.417139 | 0.395354 | 3 | 0.989692 | 0.141879 | 3 |
|  |  | SF3B1 | 1 | 0.088167 | 3 | 0.995062 | 0.26698 | 3 | 1.069934 | 0.213082 | 3 |
|  |  | SNRNP200 | 1 | 0.179631 | 3 | 0.998431 | 0.233934 | 3 | 0.921483 | 0.291165 | 3 |
|  |  | SNRNPC | 1 | 0.277926 | 3 | 0.54496 | 0.153594 | 3 | 0.475628 | 0.16707 | 3 |
| **PC3** | 3C | DDX46 | 1 | 0.326341 | 3 | 0.59184 | 0.135298 | 3 | 0.651016 | 0.079354 | 3 |
|  |  | hnRNPA1 | 1 | 0.144314 | 3 | 0.456236 | 0.135945 | 3 | 0.505155 | 0.104228 | 3 |
|  |  | hnRNPK | 1 | 0.210826 | 3 | 0.880421 | 0.089519 | 3 | 0.431863 | 0.046397 | 3 |
|  |  | HSPA8 | 1 | 0.367206 | 3 | 0.582994 | 0.17782 | 3 | 0.559089 | 0.27866 | 3 |
|  |  | SF3B1 | 1 | 0.140798 | 3 | 0.840803 | 0.142582 | 3 | 0.73925 | 0.320677 | 3 |
|  |  | SNRNP200 | 1 | 0.192376 | 3 | 0.694328 | 0.137696 | 3 | 0.421733 | 0.138391 | 3 |
|  |  | SNRNPC | 1 | 0.03748 | 3 | 0.81525 | 0.238845 | 3 | 0.822214 | 0.217901 | 3 |
| **DU145** | 3D | DDX46 | 1 | 0.133046 | 3 | 0.86597 | 0.104515 | 3 | 0.627229 | 0.052649 | 3 |
|  |  | hnRNPA1 | 1 | 0.165565 | 3 | 1.068333 | 0.136598 | 3 | 1.101365 | 0.459169 | 3 |
|  |  | hnRNPK | 1 | 0.142229 | 3 | 0.861914 | 0.11475 | 3 | 0.485053 | 0.147898 | 3 |
|  |  | HSPA8 | 1 | 0.100409 | 3 | 1.024315 | 0.174162 | 3 | 0.852377 | 0.2506 | 3 |
|  |  | SF3B1 | 1 | 0.10708 | 3 | 0.901495 | 0.041491 | 3 | 0.592128 | 0.170477 | 3 |
|  |  | SNRNP200 | 1 | 0.177283 | 3 | 0.808367 | 0.068763 | 3 | 0.594419 | 0.092736 | 3 |
|  |  | SNRNPC | 1 | 0.483977 | 3 | 1.118727 | 0.388724 | 3 | 0.640958 | 0.178394 | 3 |
|  |  |  | **VO** | | | **FOXA1** | | |  |  |  |
| **Cell Line** | **Figure** |  | **Mean** | **STDEV** | **N** | **Mean** | **STDEV** | **N** |  |  |  |
| **PC3** | 3C | DDX46 | 1 | 0.314126 | 3 | 2.31 | 1.13327 | 3 |  |  |  |
|  |  | HNRNPA1 | 1 | 0.377322 | 3 | 2.496702 | 0.966448 | 3 |  |  |  |
|  |  | HNRNPK | 1 | 0.383718 | 3 | 1.621957 | 0.063137 | 3 |  |  |  |
|  |  | HSPA8 | 1 | 0.38098 | 3 | 1.93132 | 0.851454 | 3 |  |  |  |
|  |  | SF3B1 | 1 | 0.059877 | 3 | 1.256816 | 0.03222 | 3 |  |  |  |
|  |  | SNRNP200 | 1 | 0.264982 | 3 | 2.692896 | 0.28437 | 3 |  |  |  |
|  |  | SNRNPC | 1 | 0.424927 | 3 | 1.308231 | 0.705458 | 3 |  |  |  |
|  |  |  | **NSI** | | | **siERG** | | |  |  |  |
| **Cell Line** | **Figure** | **Gene** | **Mean** | **STDEV** | **N** | **Mean** | **STDEV** | **N** |  |  |  |
| **VCaP** | S3A | DDX46 | 1 | 0.114294 | 3 | 1.326598 | 0.091924 | 3 |  |  |  |
|  |  | hnRNPA1 | 1 | 0.379305 | 3 | 0.955301 | 0.062788 | 3 |  |  |  |
|  |  | hnRNPK | 1 | 0.353075 | 3 | 1.255943 | 0.365227 | 3 |  |  |  |
|  |  | HSPA8 | 1 | 0.640009 | 3 | 1.41816 | 0.08866 | 3 |  |  |  |
|  |  | SF3B1 | 1 | 0.137483 | 3 | 1.055925 | 0.055074 | 3 |  |  |  |
|  |  | SNRNP200 | 1 | 0.27083 | 3 | 1.905663 | 0.41963 | 3 |  |  |  |
|  |  | SNRNPC | 1 | 0.526817 | 3 | 1.799998 | 0.436888 | 3 |  |  |  |
|  |  |  | **NSI** | | | **siAR** | | |  |  |  |
|  | **Figure** | **Gene** | **Mean** | **STDEV** | **N** | **Mean** | **STDEV** | **N** |  |  |  |
| **VCaP** | S3B | DDX46 | 1 | 0.114294 | 3 | 1.326598 | 0.091924 | 3 |  |  |  |
|  |  | hnRNPA1 | 1 | 0.379305 | 3 | 0.955301 | 0.062788 | 3 |  |  |  |
|  |  | hnRNPK | 1 | 0.353075 | 3 | 1.255943 | 0.365227 | 3 |  |  |  |
|  |  | HSPA8 | 1 | 0.640009 | 3 | 1.41816 | 0.08866 | 3 |  |  |  |
|  |  | SF3B1 | 1 | 0.137483 | 3 | 1.055925 | 0.055074 | 3 |  |  |  |
|  |  | SNRNP200 | 1 | 0.27083 | 3 | 1.905663 | 0.41963 | 3 |  |  |  |
|  |  | SNRNPC | 1 | 0.526817 | 3 | 1.799998 | 0.436888 | 3 |  |  |  |
| **LNCaP** | S3C | DDX46 | 1 | 0.100528 | 3 | 0.95628 | 0.225397 | 3 |  |  |  |
|  |  | hnRNPA1 | 1 | 0.199998 | 3 | 0.841509 | 0.146872 | 3 |  |  |  |
|  |  | hnRNPK | 1 | 0.184363 | 3 | 0.899902 | 0.098459 | 3 |  |  |  |
|  |  | HSPA8 | 1 | 0.382258 | 3 | 1.170482 | 0.307627 | 3 |  |  |  |
|  |  | SF3B1 | 1 | 0.042691 | 3 | 0.884945 | 0.188746 | 3 |  |  |  |
|  |  | SNRNP200 | 1 | 0.181053 | 3 | 0.732612 | 0.195074 | 3 |  |  |  |
|  |  | SNRNPC | 1 | 0.134584 | 3 | 0.851459 | 0.240133 | 3 |  |  |  |

Supplementary Table 7. Statistical Test Results Results of one-tailed Student T test between gene targeting siRNAs and NSI control, or between vector only and FOXA1 cDNA expression

| **Figure panel** | **Cell line** | **Gene** | **Controls** | **Cases** | **P value** | **Alternative** | **t, df** | **Mean ± SEM of Controls** | **Mean ± SEM of Cases** | ** means** | **95% C.I.** |
| --- | --- | --- | --- | --- | --- | --- | --- | --- | --- | --- | --- |
| 3A | VCaP | DDX46 | NSI | siFOXA1 1 | 0.1015 | One-tailed | t=1.387 df=8 | 1.000 ± 0.2484 N=5 | 0.6415 ± 0.07181 N=5 | 0.3585 ± 0.2586 | -0.2377 to 0.9548 |
|  |  |  | NSI | siFOXA1 2 | 0.0187 | One-tailed | t=2.493 df=8 | 1.000 ± 0.2484 N=5 | 0.3753 ± 0.03337 N=5 | 0.6247 ± 0.2506 | 0.04674 to 1.203 |
|  |  | HNRNPA1 | NSI | siFOXA1 1 | 0.0058 | One-tailed | t=3.258 df=8 | 1.000 ± 0.05724 N=5 | 0.7496 ± 0.05127 N=5 | 0.2504 ± 0.07684 | 0.07318 to 0.4276 |
|  |  |  | NSI | siFOXA1 2 | <0.0001 | One-tailed | t=6.511 df=8 | 1.000 ± 0.05724 N=5 | 0.5443 ± 0.04028 N=5 | 0.4557 ± 0.06999 | 0.2943 to 0.6171 |
|  |  | HNRNPK | NSI | siFOXA1 1 | 0.0978 | One-tailed | t=1.412 df=8 | 1.000 ± 0.08658 N=5 | 0.7141 ± 0.1830 N=5 | 0.2859 ± 0.2024 | -0.1809 to 0.7527 |
|  |  |  | NSI | siFOXA1 2 | 0.0423 | One-tailed | t=1.968 df=8 | 1.000 ± 0.08658 N=5 | 0.6902 ± 0.1315 N=5 | 0.3098 ± 0.1574 | -0.05322 to 0.6729 |
|  |  | HSPA8 | NSI | siFOXA1 1 | 0.0949 | One-tailed | t=1.433 df=8 | 1.000 ± 0.1213 N=5 | 0.8091 ± 0.05523 N=5 | 0.1909 ± 0.1333 | -0.1164 to 0.4982 |
|  |  |  | NSI | siFOXA1 2 | 0.0045 | One-tailed | t=3.422 df=8 | 1.000 ± 0.1213 N=5 | 0.5646 ± 0.03848 N=5 | 0.4354 ± 0.1272 | 0.1420 to 0.7288 |
|  |  | SF3B1 | NSI | siFOXA1 1 | 0.0044 | One-tailed | t=3.437 df=8 | 1.000 ± 0.05872 N=5 | 0.7930 ± 0.01346 N=5 | 0.2070 ± 0.06025 | 0.06812 to 0.3460 |
|  |  |  | NSI | siFOXA1 2 | 0.0843 | One-tailed | t=1.513 df=8 | 1.000 ± 0.05872 N=5 | 0.8691 ± 0.06346 N=5 | 0.1309 ± 0.08646 | -0.06853 to 0.3302 |
|  |  | SNRNP200 | NSI | siFOXA1 1 | 0.0046 | One-tailed | t=3.407 df=8 | 1.000 ± 0.05147 N=5 | 0.7071 ± 0.06884 N=5 | 0.2929 ± 0.08596 | 0.09467 to 0.4911 |
|  |  |  | NSI | siFOXA1 2 | <0.0001 | One-tailed | t=9.784 df=8 | 1.000 ± 0.05147 N=5 | 0.4017 ± 0.03301 N=5 | 0.5983 ± 0.06115 | 0.4573 to 0.7393 |
|  |  | SNRNPC | NSI | siFOXA1 1 | 0.1816 | One-tailed | t=0.9642 df=8 | 1.000 ± 0.3520 N=5 | 0.5661 ± 0.2803 N=5 | 0.4339 ± 0.4500 | -0.6038 to 1.472 |
|  |  |  | NSI | siFOXA1 2 | 0.0849 | One-tailed | t=1.509 df=8 | 1.000 ± 0.3520 N=5 | 0.3986 ± 0.1868 N=5 | 0.6014 ± 0.3985 | -0.3176 to 1.520 |
| 3B | LNCaP | DDX46 | NSI | siFOXA1 1 | 0.0694 | One-tailed | t=1.845 df=4 | 1.000 ± 0.08414 N=3 | 1.380 ± 0.1878 N=3 | -0.3795 ± 0.2058 | -0.9507 to 0.1917 |
|  |  |  | NSI | siFOXA1 2 | 0.4109 | One-tailed | t=0.2405 df=4 | 1.000 ± 0.08414 N=3 | 1.037 ± 0.1300 N=3 | -0.03724 ± 0.1548 | -0.4671 to 0.3926 |
|  |  | HNRNPA1 | NSI | siFOXA1 1 | 0.1463 | One-tailed | t=1.211 df=4 | 1.000 ± 0.1045 N=3 | 0.8497 ± 0.06701 N=3 | 0.1503 ± 0.1242 | -0.1943 to 0.4950 |
|  |  |  | NSI | siFOXA1 2 | 0.0015 | One-tailed | t=6.455 df=4 | 1.000 ± 0.1045 N=3 | 0.3002 ± 0.02876 N=3 | 0.6998 ± 0.1084 | 0.3988 to 1.001 |
|  |  | HNRNPK | NSI | siFOXA1 1 | 0.3891 | One-tailed | t=0.3012 df=4 | 1.000 ± 0.09750 N=3 | 0.9662 ± 0.05571 N=3 | 0.03382 ± 0.1123 | -0.2779 to 0.3455 |
|  |  |  | NSI | siFOXA1 2 | 0.0068 | One-tailed | t=4.210 df=4 | 1.000 ± 0.09750 N=3 | 0.5016 ± 0.06713 N=3 | 0.4984 ± 0.1184 | 0.1698 to 0.8270 |
|  |  | HSPA8 | NSI | siFOXA1 1 | 0.0914 | One-tailed | t=1.610 df=4 | 1.000 ± 0.1227 N=3 | 1.417 ± 0.2283 N=3 | -0.4171 ± 0.2592 | -1.137 to 0.3023 |
|  |  |  | NSI | siFOXA1 2 | 0.4738 | One-tailed | t=0.06987 df=4 | 1.000 ± 0.1227 N=3 | 0.9897 ± 0.08191 N=3 | 0.01031 ± 0.1475 | -0.3993 to 0.4199 |
|  |  | SF3B1 | NSI | siFOXA1 1 | 0.4886 | One-tailed | t=0.03042 df=4 | 1.000 ± 0.05090 N=3 | 0.9951 ± 0.1541 N=3 | 0.004938 ± 0.1623 | -0.4457 to 0.4556 |
|  |  |  | NSI | siFOXA1 2 | 0.3136 | One-tailed | t=0.5253 df=4 | 1.000 ± 0.05090 N=3 | 1.070 ± 0.1230 N=3 | -0.06993 ± 0.1331 | -0.4395 to 0.2997 |
|  |  | SNRNP200 | NSI | siFOXA1 1 | 0.4965 | One-tailed | t=0.009215 df=4 | 1.000 ± 0.1037 N=3 | 0.9984 ± 0.1351 N=3 | 0.001569 ± 0.1703 | -0.4711 to 0.4743 |
|  |  |  | NSI | siFOXA1 2 | 0.3556 | One-tailed | t=0.3975 df=4 | 1.000 ± 0.1037 N=3 | 0.9215 ± 0.1681 N=3 | 0.07852 ± 0.1975 | -0.4698 to 0.6268 |
|  |  | SNRNPC | NSI | siFOXA1 1 | 0.034 | One-tailed | t=2.482 df=4 | 1.000 ± 0.1605 N=3 | 0.5450 ± 0.08868 N=3 | 0.4550 ± 0.1833 | -0.05389 to 0.9640 |
|  |  |  | NSI | siFOXA1 2 | 0.0244 | One-tailed | t=2.801 df=4 | 1.000 ± 0.1605 N=3 | 0.4756 ± 0.09646 N=3 | 0.5244 ± 0.1872 | 0.004646 to 1.044 |
| 3C | PC3 | DDX46 | NSI | siFOXA1 1 | 0.058 | One-tailed | t=2.001 df=4 | 1.000 ± 0.1884 N=3 | 0.5918 ± 0.07811 N=3 | 0.4082 ± 0.2040 | -0.1580 to 0.9744 |
|  |  |  | NSI | siFOXA1 2 | 0.0731 | One-tailed | t=1.800 df=4 | 1.000 ± 0.1884 N=3 | 0.6510 ± 0.04582 N=3 | 0.3490 ± 0.1939 | -0.1893 to 0.8873 |
|  |  | HNRNPA1 | NSI | siFOXA1 1 | 0.0045 | One-tailed | t=4.750 df=4 | 1.000 ± 0.08332 N=3 | 0.4562 ± 0.07849 N=3 | 0.5438 ± 0.1145 | 0.2260 to 0.8615 |
|  |  |  | NSI | siFOXA1 2 | 0.0043 | One-tailed | t=4.815 df=4 | 1.000 ± 0.08332 N=3 | 0.5052 ± 0.06018 N=3 | 0.4948 ± 0.1028 | 0.2095 to 0.7802 |
|  |  | HNRNPK | NSI | siFOXA1 1 | 0.2085 | One-tailed | t=0.9043 df=4 | 1.000 ± 0.1217 N=3 | 0.8804 ± 0.05168 N=3 | 0.1196 ± 0.1322 | -0.2475 to 0.4867 |
|  |  |  | NSI | siFOXA1 2 | 0.0052 | One-tailed | t=4.558 df=4 | 1.000 ± 0.1217 N=3 | 0.4319 ± 0.02679 N=3 | 0.5681 ± 0.1246 | 0.2222 to 0.9141 |
|  |  | HSPA8 | NSI | siFOXA1 1 | 0.0757 | One-tailed | t=1.770 df=4 | 1.000 ± 0.2120 N=3 | 0.5830 ± 0.1027 N=3 | 0.4170 ± 0.2356 | -0.2369 to 1.071 |
|  |  |  | NSI | siFOXA1 2 | 0.0865 | One-tailed | t=1.657 df=4 | 1.000 ± 0.2120 N=3 | 0.5591 ± 0.1609 N=3 | 0.4409 ± 0.2661 | -0.2979 to 1.180 |
|  |  | SF3B1 | NSI | siFOXA1 1 | 0.1204 | One-tailed | t=1.376 df=4 | 1.000 ± 0.08129 N=3 | 0.7393 ± 0.1851 N=3 | 0.2608 ± 0.2022 | -0.3006 to 0.8221 |
|  |  |  | NSI | siFOXA1 2 | 0.1334 | One-tailed | t=1.290 df=4 | 1.000 ± 0.08129 N=3 | 0.8408 ± 0.08232 N=3 | 0.1592 ± 0.1157 | -0.1620 to 0.4804 |
|  |  | SNRNP200 | NSI | siFOXA1 1 | 0.0444 | One-tailed | t=2.238 df=4 | 1.000 ± 0.1111 N=3 | 0.6943 ± 0.07950 N=3 | 0.3057 ± 0.1366 | -0.07350 to 0.6848 |
|  |  |  | NSI | siFOXA1 2 | 0.0067 | One-tailed | t=4.226 df=4 | 1.000 ± 0.1111 N=3 | 0.4217 ± 0.07990 N=3 | 0.5783 ± 0.1368 | 0.1984 to 0.9581 |
|  |  | SNRNPC | NSI | siFOXA1 1 | 0.1281 | One-tailed | t=1.324 df=4 | 1.000 ± 0.02164 N=3 | 0.8152 ± 0.1379 N=3 | 0.1848 ± 0.1396 | -0.2027 to 0.5722 |
|  |  |  | NSI | siFOXA1 2 | 0.1181 | One-tailed | t=1.393 df=4 | 1.000 ± 0.02164 N=3 | 0.8222 ± 0.1258 N=3 | 0.1778 ± 0.1277 | -0.1766 to 0.5321 |
| 3D | DU145 | DDX46 | NSI | siFOXA1 1 | 0.121 | One-tailed | t=1.372 df=4 | 1.000 ± 0.07681 N=3 | 0.8660 ± 0.06034 N=3 | 0.1340 ± 0.09768 | -0.1371 to 0.4052 |
|  |  |  | NSI | siFOXA1 2 | 0.0054 | One-tailed | t=4.512 df=4 | 1.000 ± 0.07681 N=3 | 0.6272 ± 0.03040 N=3 | 0.3728 ± 0.08261 | 0.1434 to 0.6021 |
|  |  | HNRNPA1 | NSI | siFOXA1 1 | 0.3054 | One-tailed | t=0.5514 df=4 | 1.000 ± 0.09559 N=3 | 1.068 ± 0.07886 N=3 | -0.06833 ± 0.1239 | -0.4123 to 0.2757 |
|  |  |  | NSI | siFOXA1 2 | 0.3686 | One-tailed | t=0.3597 df=4 | 1.000 ± 0.09559 N=3 | 1.101 ± 0.2651 N=3 | -0.1014 ± 0.2818 | -0.8837 to 0.6809 |
|  |  | HNRNPK | NSI | siFOXA1 1 | 0.1304 | One-tailed | t=1.309 df=4 | 1.000 ± 0.08212 N=3 | 0.8619 ± 0.06625 N=3 | 0.1381 ± 0.1055 | -0.1548 to 0.4310 |
|  |  |  | NSI | siFOXA1 2 | 0.0061 | One-tailed | t=4.347 df=4 | 1.000 ± 0.08212 N=3 | 0.4851 ± 0.08539 N=3 | 0.5149 ± 0.1185 | 0.1861 to 0.8438 |
|  |  | HSPA8 | NSI | siFOXA1 1 | 0.4222 | One-tailed | t=0.2095 df=4 | 1.000 ± 0.05797 N=3 | 1.024 ± 0.1006 N=3 | -0.02432 ± 0.1161 | -0.3465 to 0.2979 |
|  |  |  | NSI | siFOXA1 2 | 0.1986 | One-tailed | t=0.9471 df=4 | 1.000 ± 0.05797 N=3 | 0.8524 ± 0.1447 N=3 | 0.1476 ± 0.1559 | -0.2851 to 0.5803 |
|  |  | SF3B1 | NSI | siFOXA1 1 | 0.1058 | One-tailed | t=1.486 df=4 | 1.000 ± 0.06182 N=3 | 0.9015 ± 0.02395 N=3 | 0.09851 ± 0.06630 | -0.08555 to 0.2826 |
|  |  |  | NSI | siFOXA1 2 | 0.0123 | One-tailed | t=3.509 df=4 | 1.000 ± 0.06182 N=3 | 0.5921 ± 0.09842 N=3 | 0.4079 ± 0.1162 | 0.08522 to 0.7305 |
|  |  | SNRNP200 | NSI | siFOXA1 1 | 0.0779 | One-tailed | t=1.746 df=4 | 1.000 ± 0.1024 N=3 | 0.8084 ± 0.03970 N=3 | 0.1916 ± 0.1098 | -0.1131 to 0.4964 |
|  |  |  | NSI | siFOXA1 2 | 0.0123 | One-tailed | t=3.511 df=4 | 1.000 ± 0.1024 N=3 | 0.5944 ± 0.05354 N=3 | 0.4056 ± 0.1155 | 0.08492 to 0.7262 |
|  |  | SNRNPC | NSI | siFOXA1 1 | 0.3785 | One-tailed | t=0.3313 df=4 | 1.000 ± 0.2794 N=3 | 1.119 ± 0.2244 N=3 | -0.1187 ± 0.3584 | -1.114 to 0.8762 |
|  |  |  | NSI | siFOXA1 2 | 0.1472 | One-tailed | t=1.206 df=4 | 1.000 ± 0.2794 N=3 | 0.6410 ± 0.1030 N=3 | 0.3590 ± 0.2978 | -0.4677 to 1.186 |
| 3E | PC3 | DDX46 | VO | FOXA1 | 0.063 | One-tailed | t=1.929 df=4 | 1.000 ± 0.1814 N=3 | 2.310 ± 0.6543 N=3 | -1.310 ± 0.6790 | -3.195 to 0.5748 |
|  |  | HNRNPA1 | VO | FOXA1 | 0.0334 | One-tailed | t=2.499 df=4 | 1.000 ± 0.2178 N=3 | 2.497 ± 0.5580 N=3 | -1.497 ± 0.5990 | -3.160 to 0.1661 |
|  |  | HNRNPK | VO | FOXA1 | 0.0252 | One-tailed | t=2.770 df=4 | 1.000 ± 0.2215 N=3 | 1.622 ± 0.03645 N=3 | -0.6220 ± 0.2245 | -1.245 to 0.001307 |
|  |  | HSPA8 | VO | FOXA1 | 0.0794 | One-tailed | t=1.729 df=4 | 1.000 ± 0.2200 N=3 | 1.931 ± 0.4916 N=3 | -0.9313 ± 0.5386 | -2.426 to 0.5637 |
|  |  | SF3B1 | VO | FOXA1 | 0.0014 | One-tailed | t=6.542 df=4 | 1.000 ± 0.03457 N=3 | 1.257 ± 0.01860 N=3 | -0.2568 ± 0.03926 | -0.3658 to -0.1478 |
|  |  | SNRNP200 | VO | FOXA1 | 0.0008 | One-tailed | t=7.544 df=4 | 1.000 ± 0.1530 N=3 | 2.693 ± 0.1642 N=3 | -1.693 ± 0.2244 | -2.316 to -1.070 |
|  |  | SNRNPC | VO | FOXA1 | 0.2761 | One-tailed | t=0.6483 df=4 | 1.000 ± 0.2453 N=3 | 1.308 ± 0.4073 N=3 | -0.3082 ± 0.4755 | -1.628 to 1.012 |
| S3A | VCaP | DDX46 | NSI | siERG | 0.0091 | One-tailed | t=3.857 df=4 | 1.000 ± 0.06599 N=3 | 1.327 ± 0.05307 N=3 | -0.3266 ± 0.08468 | -0.5617 to -0.09152 |
|  |  | HNRNPA1 | NSI | siERG | 0.4251 | One-tailed | t=0.2014 df=4 | 1.000 ± 0.2190 N=3 | 0.9553 ± 0.03625 N=3 | 0.04470 ± 0.2220 | -0.5715 to 0.6609 |
|  |  | HNRNPK | NSI | siERG | 0.216 | One-tailed | t=0.8727 df=4 | 1.000 ± 0.2038 N=3 | 1.256 ± 0.2109 N=3 | -0.2559 ± 0.2933 | -1.070 to 0.5582 |
|  |  | HSPA8 | NSI | siERG | 0.1625 | One-tailed | t=1.121 df=4 | 1.000 ± 0.3695 N=3 | 1.418 ± 0.05119 N=3 | -0.4182 ± 0.3730 | -1.454 to 0.6174 |
|  |  | SF3B1 | NSI | siERG | 0.2744 | One-tailed | t=0.6540 df=4 | 1.000 ± 0.07938 N=3 | 1.056 ± 0.03180 N=3 | -0.05593 ± 0.08551 | -0.2933 to 0.1814 |
|  |  | SNRNP200 | NSI | siERG | 0.0174 | One-tailed | t=3.141 df=4 | 1.000 ± 0.1564 N=3 | 1.906 ± 0.2423 N=3 | -0.9057 ± 0.2884 | -1.706 to -0.1052 |
|  |  | SNRNPC | NSI | siERG | 0.0565 | One-tailed | t=2.025 df=4 | 1.000 ± 0.3042 N=3 | 1.800 ± 0.2522 N=3 | -0.8000 ± 0.3951 | -1.897 to 0.2969 |
| S3B | VCaP | DDX46 | NSI | siAR | 0.1345 | One-tailed | t=1.282 df=4 | 1.000 ± 0.2115 N=3 | 1.290 ± 0.08081 N=3 | -0.2903 ± 0.2264 | -0.9188 to 0.3382 |
|  |  | HNRNPA1 | NSI | siAR | 0.2025 | One-tailed | t=0.9299 df=4 | 1.000 ± 0.1971 N=3 | 0.7692 ± 0.1509 N=3 | 0.2308 ± 0.2482 | -0.4582 to 0.9198 |
|  |  | HNRNPK | NSI | siAR | 0.214 | One-tailed | t=0.8810 df=4 | 1.000 ± 0.2953 N=3 | 1.278 ± 0.1114 N=3 | -0.2780 ± 0.3156 | -1.154 to 0.5980 |
|  |  | HSPA8 | NSI | siAR | 0.1417 | One-tailed | t=1.238 df=4 | 1.000 ± 0.2444 N=3 | 0.6698 ± 0.1070 N=3 | 0.3302 ± 0.2668 | -0.4104 to 1.071 |
|  |  | SF3B1 | NSI | siERG | 0.1272 | One-tailed | t=1.330 df=4 | 1.000 ± 0.1162 N=3 | 1.218 ± 0.1154 N=3 | -0.2178 ± 0.1638 | -0.6725 to 0.2369 |
|  |  | SNRNP200 | NSI | siAR | 0.3825 | One-tailed | t=0.3200 df=4 | 1.000 ± 0.2464 N=3 | 1.100 ± 0.1916 N=3 | -0.09986 ± 0.3121 | -0.9662 to 0.7665 |
|  |  | SNRNPC | NSI | siAR | 0.0928 | One-tailed | t=1.596 df=4 | 1.000 ± 0.3165 N=3 | 1.619 ± 0.2244 N=3 | -0.6193 ± 0.3880 | -1.696 to 0.4578 |
| S3C | LNCaP | DDX46 | NSI | siAR | 0.3871 | One-tailed | t=0.3068 df=4 | 1.000 ± 0.05804 N=3 | 0.9563 ± 0.1301 N=3 | 0.04372 ± 0.1425 | -0.3518 to 0.4393 |
|  |  | HNRNPA1 | NSI | siAR | 0.1653 | One-tailed | t=1.106 df=4 | 1.000 ± 0.1155 N=3 | 0.8415 ± 0.08480 N=3 | 0.1585 ± 0.1433 | -0.2392 to 0.5562 |
|  |  | HNRNPK | NSI | siAR | 0.2267 | One-tailed | t=0.8295 df=4 | 1.000 ± 0.1064 N=3 | 0.8999 ± 0.05685 N=3 | 0.1001 ± 0.1207 | -0.2349 to 0.4351 |
|  |  | HSPA8 | NSI | siAR | 0.2899 | One-tailed | t=0.6018 df=4 | 1.000 ± 0.2207 N=3 | 1.170 ± 0.1776 N=3 | -0.1705 ± 0.2833 | -0.9569 to 0.6159 |
|  |  | SF3B1 | NSI | siERG | 0.1806 | One-tailed | t=1.030 df=4 | 1.000 ± 0.02465 N=3 | 0.8849 ± 0.1090 N=3 | 0.1151 ± 0.1117 | -0.1951 to 0.4252 |
|  |  | SNRNP200 | NSI | siAR | 0.0784 | One-tailed | t=1.740 df=4 | 1.000 ± 0.1045 N=3 | 0.7326 ± 0.1126 N=3 | 0.2674 ± 0.1537 | -0.1592 to 0.6939 |
|  |  | SNRNPC | NSI | siAR | 0.2014 | One-tailed | t=0.9346 df=4 | 1.000 ± 0.07770 N=3 | 0.8515 ± 0.1386 N=3 | 0.1485 ± 0.1589 | -0.2927 to 0.5897 |

Results of one-tailed Student T test between gene targeting siRNAs and NSI control

Supplementary Table 8. Sequences and calculated efficiencies for RT-qPCR primers used in this study

| **Gene** | **Species** | **Forward Primer** | **Reverse Primer** | **UPL Probe** |
| --- | --- | --- | --- | --- |
| ACTB | Human | agagctacgagctgcctgac | cgtggatgccacaggact | 9 |
| AR | Human | ggactccgtgcagcctatt | gggcacttgcacagagatg | 53 |
| B2M | Human | ttctggcctggaggctatc | tcaggaaatttgactttccattc | 42 |
| DDX46 | Human | aatactctgaagccgcaattaca | cagccagttcattggcact | 6 |
| ERG | Human | aagtagccgccttgcaaat | tgatgcagctggagttgg | *SYBR* |
| FOXA1 | Human | agggctggatggttgtattg | accgggacggaggagtag | 1 |
| hnRNPA1 | Human | gtggggatggctataatgga | tccatagcctctgcttcctc | 25 |
| hnRNPK | Human | gcgctcgttttctgtctagc | cccagtgctgcagtagcc | 30 |
| HOXB13 | Human | ttttggaaggcagcatttg | gaacttgttagccgcatactcc | 24 |
| HSPA8 | Human | tgctgctgctattgcttacg | ctcaaagattccatcctcaa | 10 |
| SF3B1 | Human | aaggccattgtaaatgtcatagg | tgttctttaagatgggggtga | *SYBR* |
| SNRNP200 | Human | ttctatgatgatgccatcgtgt | ctggccgtcttcaaaatctc | 8 |
| SNRNPC | Human | ctgcaggggcgatgatac | caggagcaggtcccactg | 79 |

Additional data table S1 (separate file)

TCGA PRAD Patient ID and expression information for five transcription factors

Additional data table S2 (separate file)

Expression of SRPs in *FOXA1* HE versus REST, DETEMER Scores for SRP dependency from Project Achilles data and ReMap analysis of TF binding sites within SRP gene loci. Expression of SRPs in *FOXA1* HE versus REST for SU2C dataset.
